# Mutations of Omicron variant at the interface of the receptor domain motif and human angiotensin-converting enzyme-2

**DOI:** 10.1101/2022.02.04.479136

**Authors:** Puja Adhikari, Bahaa Jawad, Rudolf Podgornik, Wai-Yim Ching

**Author notes:** Correspondence: Wai-Yim Ching.

## Abstract

The most recent Omicron variant of SARS-CoV-2 has caused global concern and anxiety. The only thing certain about this strain, with large number of mutations in the spike protein, is that it spreads quickly, seems to evade immune defense and mitigates the benefits of existing vaccines. Based on the ultra-large-scale *ab initio* computational modeling of the receptor binding motif (RBM) and human angiotensin-converting enzyme-2 (ACE2) interface we provide the details of the effect of Omicron mutations at the fundamental atomic scale level. In-depth analysis anchored in the novel concept of *amino acid-amino acid bond pair units* (AABPU), indicates that mutations in the Omicron variant are connected with (i) significant changes in the shape and structure of AABPU components, together with (ii) significant increase in the positive partial charge which facilitates the interaction with ACE2. The calculated bond order, based on AABPU, reveals that the Omicron mutations increase the binding strength of RBM to ACE2. Our findings correlate with and are instrumental to explain the current observations and can contribute to the prediction of next potential new variant of concern.

## 1. Introduction

COVID-19 caused by SARS-CoV-2 has adversely affected global health and economics for more than two years now. It has taken millions of lives and caused chaos in every aspect of our lives. Despite significant progress in SARS-CoV-2 research yielding effective vaccines, the virus continues to mutate and develop, providing ever new variants of concern (VOCs) [1-5] and variants of interests (VOIs) [6-10] that have instigated new anxieties. The newly emerging variants can change virus properties such as transmissibility, antigenicity, infectivity, and pathogenicity [11]. The Omicron variant (OV) is the most recently identified SARS-CoV-2 VOCs that has a substantially larger number of mutations and a much higher rate of transmission than previous variants [12-14]. The first case of OV was reported to WHO from South Africa on 24 November 2021 [15]. However, ever since the OV has been rapidly spreading all over the world and has become the dominant variant. In fact, OV has been estimated to be 99.5% of all cases in US by Centers for Disease Control and Prevention (CDC) [16]. The transmissibility and severity of this variant, as well as its ability to evade vaccines and causing reinfections, remains unknown. These developments instigated a strong research focus to identify the OV impact on the efficacy of vaccines and to improve/develop drugs to mitigate its effects. Hence, elucidating the effects of OV mutations on the binding with human cell is necessary to provide solid understanding of the molecular basis of this variant.

OV has more than 30 mutations in the spike (S)-protein, including 15 mutations in its receptor binding domain (RBD) [17]. The S-protein of SARS-CoV-2 is responsible for the viral entry into the human cell and activating the infection, in particular since RBD attaches directly to human angiotensin-converting enzyme 2 (ACE2).Therefore S-protein and its RBD are considered as targets for vaccination and therapeutic development [18-22]. The receptor binding motif (RBM) in the RBD is the main functional motif that forms the interface between S-protein and ACE2 [23-25]. Among the 15 mutations in RBD, RBM consists of 10 mutations: N440K, G446S, S477N, T478K, E484A, Q493R, G496S, Q498R, N501Y, Y505H [14]. This unusually large number of OV mutations has promoted valid concerns about their impact on the efficacy of existing vaccines and treatments [26-29]. More specifically, these concerns become further exacerbated when it was discovered that some OV mutations at RBM like T478K, E484A and N501Y, which been found in previous variants [30] have been demonstrated to influence ACE2 binding or have been involved in escaping antibodies [30-32]. On the other hand, the other seven mutations of RBM are unique to OV, and their biological functions are still undetermined.

In this context, the investigation of how these OV mutations interact with ACE2 human receptor is crucial for understanding the efficiency of viral entry and the ensuing speed of viral proliferation. Moreover, it could provide also important information regarding identification of significant epitopes on RBM to guide the therapeutic development against SARS-CoV-2 variants. In this work, we focus on the 10 mutations in RBM of OV and the changes they instigate, in the interaction of amino acids (AAs) at the interface of RBM and ACE2, between the unmutated or Wild Type (WT) and mutated OV virus types. *Ab initio* quantum mechanical calculation based on density functional theory (DFT) has been implemented to gain deep insights of these interaction at atomic as well as AA scale. A novel concept of *amino acid amino acid bond pair unit* (AABPU) has been developed to emphasize the changes in the bonding and other properties such as shape, volume, surface, and their partial charge.

## 2. Model construction

The present work focuses on the interactions in the interface complex between RBM and a portion of ACE2 derived from the PDB ID 6M0J [24], as detailed in our previous publication [33]. This system contains 71 amino acids (AAs) from S438 to Y508 of the RBM and 117 AAs from S19 to I88 and G319 to T365 of the ACE2. To neutralize the model, we have also added 6 Na^+^ ions and a single Na^+^ ion to unmutated or the Wild Type (WT) model and Omicron mutated model, respectively. The placing of the Na^+^ ions was performed *via* a Coulomb potential on a grid, using the LEaP program in the AMBER (Assisted Model Building with Energy Refinement) package. Hydrogen (H) atoms were added using LEaP module [34, 35]. We used the Dunbrack backbone-dependent rotamer library [36] implemented by UCSF Chimera [37] to mutate the 10 AAs for the Omicron variant. For torsion and angle adjustment of T478K and N501Y we used data from PDB ID 7ORA [8] and 7V80 [38], respectively. The WT and OV model consist of total 2930 atoms and 2964 atoms, respectively, as illustrated in **Figure 1**. These two interface models (WT and OV) are fully optimized using the Vienna *ab initio* simulation package (VASP) [39] and then the optimized structure is used as input for electronic structure calculation using orthogonalized linear combination of atomic orbitals (OLCAO) [40]. Details of these methods are discussed in **Section S1** of Supplementary Material (**SM**). The combination of using these two different DFT codes is well documented and is especially effective for large complex biomolecular systems such as the S-protein [33, 41-44].

**Figure 1.**
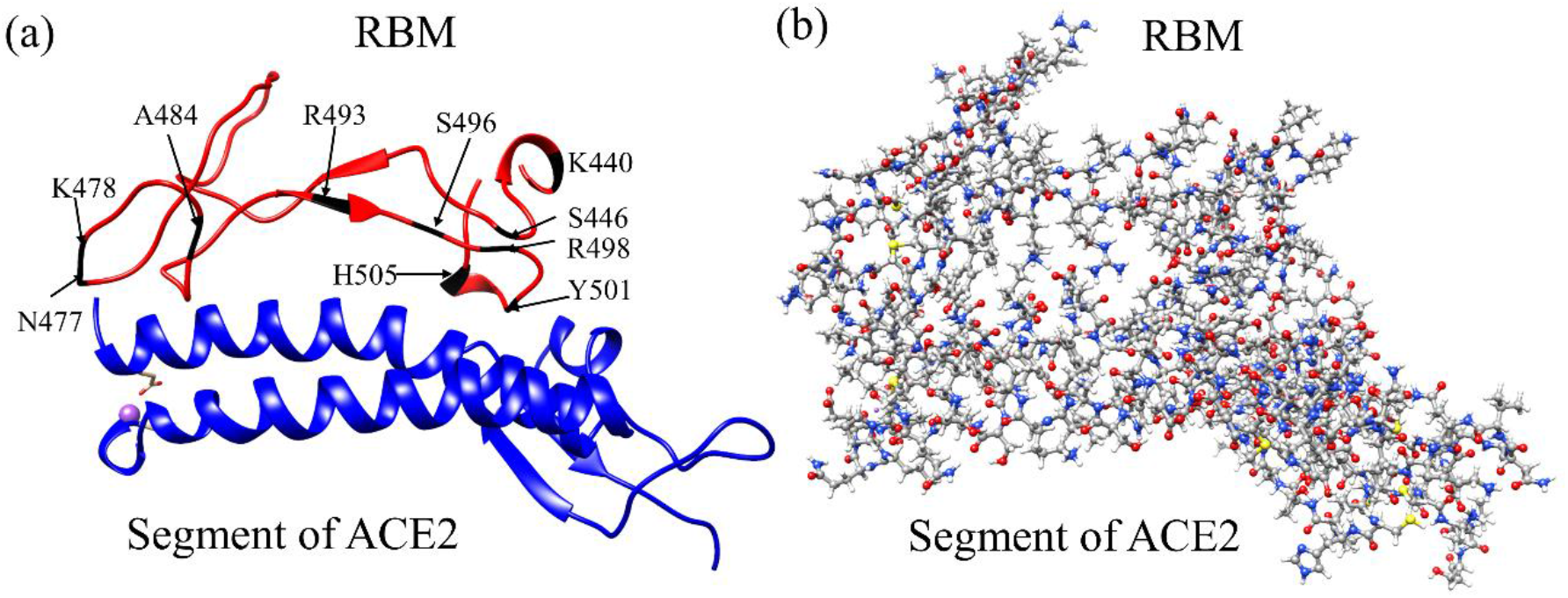
The RBM-ACE2 interface model. (a) Ribbon structure showing the interface between RBM and segment of ACE2. 10 mutated AAs of the RBM in OV are marked. (b) The ball and stick structure of the same model in (a): Grey: C, red: O, blue: N, and white: H. In the WT interface model, there are 1102 atoms in RBM, 1822 atoms in ACE2 segment, and 6 Na^+^ ions with a total of 2930 atoms. In OV interface model, there are 1141 atoms in RBM, 1822 atoms in ACE2 segment, and 1 Na^+^ ions with a total of 2964 atoms.

## 3. Result

In OLCAO, the bond order (BO) or strength of bond is calculated using Mulliken’s Scheme [45, 46]. The traditional BO description has been extended to quantify the bonding strength between two amino acids (u,*v*) called *amino acid-amino acid bond pair*, AABP(u,v) [47] in Eq.(1), since in biomolecular systems, the use of AABP is more useful than interatomic bonding between a pair of atoms.

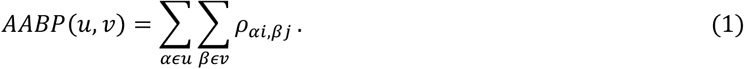

The AABP considers all possible bonding between two amino acids including both covalent and hydrogen bonding (HB). This single quantitative parameter, provided by the electronic structure, reflects the internal bonding strength between amino acids. AABP can be further resolved into nearest neighbor (NN) pairs and non-local (NL) bonding from non-NN pairs along the protein backbone. We can further identify, the contribution arising from HB to AABP. Hence AABP is an ideal parameter to characterize interactions between different AAs or groups of interacting AAs in biomolecules. In order words, we can consider AABP to be the basic biological unit that we refer to as the AAPB unit (AABPU). This will be thoughtfully elaborated in the following section.

### 3.1 Analysis of AABPU for mutation in RBM-ACE2

In this section, we characterize the overall behavior of ten mutations in RBM-ACE2 interface model for the OV in terms of the key parameters in AABPU: total AABP, NN AABP, AABP due to NL interactions, AABP from HBs, number of NL AAs, the volume and surface area of the AABPU, and the calculated partial charge (PC) for the AABPU, see **Table 1**. Pairs of subsequent rows correspond to an unmutated WT site (light blue) and a mutated OV site (light orange), respectively. For example, the first 2 rows show the calculated key parameters for N440K mutation, where in both WT and OV cases N440 and K440 interact with 2 NN Aas and 3NL AAs form two different AABPUs of 6 AAs each but with different volumes as well as values for CI and PC. The change in shape for the 10 mutations in the RBM-ACE2 model is depicted in **Figure 2**.

**Table 1.** Comparison of results between WT and OV for their AABP units in ten RBM sites. The two mutations in red have their volumes decreased after mutation.

| Models | Total AABP | NN AABP | NL AABP | AABP from HB | # NL AAs | Volume (Å<sup>3</sup>) | Area (Å<sup>2</sup>) | PC*(e<sup>-</sup>) |
| --- | --- | --- | --- | --- | --- | --- | --- | --- |
| WT N440 | 0.983 | 0.979 | 0.005 | 0.037 | 3 | 590.9 | 466.8 | -1.096 |
| OV K440 | 0.983 | 0.977 | 0.006 | 0.037 | 3 | 608.9 | 502.5 | -0.402 |
| WT G446 | 0.976 | 0.945 | 0.032 | 0.067 | 3 | 601.9 | 530.6 | 0.641 |
| OV S446 | 1.073 | 1.013 | 0.060 | 0.116 | 4 | 838.8 | 704.2 | 1.404 |
| WT S477 | 0.962 | 0.955 | 0.007 | 0.039 | 2 | 447.1 | 388.7 | 0.085 |
| OV N477 | 1.178 | 1.177 | 0.001 | 0.170 | 2 | 500.9 | 441.9 | 1.074 |
| WT T478 | 1.064 | 1.062 | 0.002 | 0.022 | 2 | 485 | 427.1 | 0.027 |
| OV K478 | 1.258 | 1.255 | 0.003 | 0.156 | 3 | 594.4 | 507.9 | 1.058 |
| WT E484 | 1.049 | 0.930 | 0.119 | 0.133 | 4 | 823 | 678.8 | -0.411 |
| OV A484 | 0.947 | 0.944 | 0.002 | 0.032 | 2 | 447.9 | 413.5 | -0.142 |
| WT Q493 | 1.197 | 0.969 | 0.229 | 0.241 | 8 | 1482 | 986.5 | 0.188 |
| OV R493 | 1.263 | 1.058 | 0.205 | 0.270 | 8 | 1467 | 1070 | 0.644 |
| WT G496 | 1.127 | 0.992 | 0.135 | 0.158 | 7 | 1247 | 929.2 | -0.021 |
| OV S496 | 1.098 | 0.964 | 0.134 | 0.146 | 7 | 1359 | 952.2 | 0.874 |
| WT Q498 | 1.117 | 1.072 | 0.045 | 0.052 | 11 | 1502 | 1019 | 0.926 |
| OV R498 | 1.220 | 1.058 | 0.162 | 0.147 | 15 | 2346 | 1365 | 1.889 |
| WT N501 | 1.091 | 0.963 | 0.128 | 0.135 | 8 | 1251 | 883.8 | 0.447 |
| OV Y501 | 1.139 | 0.957 | 0.183 | 0.135 | 9 | 1591 | 988.9 | 0.536 |
| WT Y505 | 1.038 | 0.960 | 0.078 | 0.102 | 5 | 847.2 | 637.8 | 0.460 |
| OV H505 | 1.020 | 0.934 | 0.086 | 0.107 | 6 | 993.5 | 720.4 | -0.188 |

**Figure 2.**
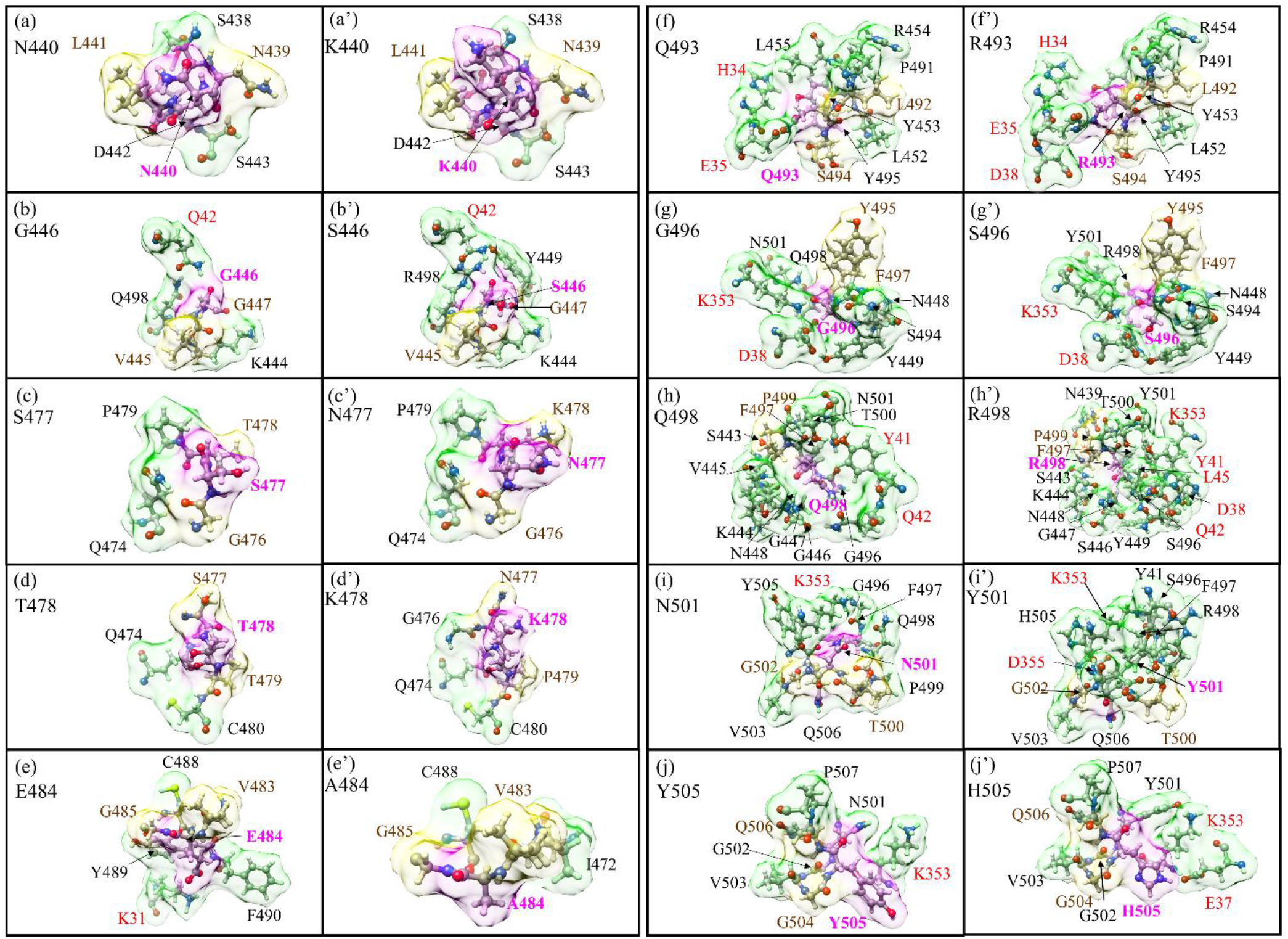
Details of the shape change of AABPU of the ten mutation sites in RBM: (a) N440, (b) G446, (c) S477, (d) T478, (e) E484, (f) Q493, (g) G496, (h) Q498, (i) N501, and (j) Y505 for the WT. (a’) K440, (b’) S446, (c’) N477, (d’) K478, (e’) A484, (f’) R493, (g’) S496, (h’) R498, (i’) Y501, and (j’) H505 for the OV. The surface of mutated sites is shown in magenta, surface of NN and NL are shown in yellow and green respectively. All NN and NL AAs are marked near to their surface in brown and black respectively. NL AAs from ACE2 are marked in red.

In **Figure. 2**, we show the change in shape of the 10 AABPU for WT and OV, side by side, for easy comparison, with the details of bonding configurations not shown in **Figure 2** listed in **Table S1** to **Table S20** in **SI** for WT and OV shown side by side. In these tables, the colors for the NN, NL bonds are in yellow and green, consistent with the color used for surface shown in **Figure 2**. Detailed inspection of **Figure 2** reveals substantial changes in shapes, orientations, and number of NL AAs due to mutations. The most prominent is exhibited by the Q498 to R498 mutation, where the number of NL AAs has increased from 11 to 15 (**Figure 2 (h)** and **(h’)**). On the other hand, for the mutation from E484 to A484 the number of NL AAs decreased from 4 to 2 (**Figure 2 (e)** and **(e’)**). These changes will affect the overall bonding in the AABPU of the mutated sites.

The most prominent mutation effect is exhibited by Q498 to R498 mutation, where the number of NL AAs has increased from 11 to 15. The AABPU, as mentioned above, is an important aggregation of parameters to show the changes in the interaction due to mutation. There are several important points from **Table 1** and **Figure S1** that should be explicitly pointed out. (1) Mutation increases the total AABP in 6 out of 10 sites. i.e., G446S, S477N, T478K, Q493R, Q498R, and N501Y. (2) The change in total AABP is affected by both NN AABP and NL AABP. Specifically, mutation increases the NN AABP in 5 out of 10 sites (G446S, S477N, T478K, E484A, and Q493R), and increases the NL AABP in 5 out of 10 sites (G446S, T478K, Q498R, N501Y, Y505H). Hence mutation increases or decreases the overall bonding strength depending on the site of mutation. Each site is unique in terms of inter-amino acid interaction and behaves accordingly. (3) Mutation increases the contribution from HB to total AABP in 6 out of 10 sites. (G446S, S477N, T478K, Q493R, Q498R, and Y505H). (4) Mutation increases the number of NL AAs in 5 out of 10 sites (G446S, T478K, Q498R, N501Y, Y505H) but just by one. However, in site Q498R the NL AAs increases by four and in E484A decreases by two. In fact, E484A is the only site where the number of interacting NL AAs decreases after mutation. (5) Increase in number of NL AAs seems to increase its total AABP. However, there could be outlier such as in Y505H which has one more NL AA but its total AABP gets lower after mutation due to decrease in the NN AABP after mutation.

In analyzing AABPU we focus on the change in volume and surface area of the unit. Mutation increases the volume in 8 out of 10 sites except in E484A and Q493R. In E484A, the number of interacting NL AAs decreases from 4 to 2, the volume decreases by 45.6% and the surface area by 39.1%. In Q493R, The NL AAs are the same (8), the decrease in volume (1.0%) is negligible and the increase in surface area (19.4%) is relatively small. On the other hand, the most prominent changes occur in Q498R the NL AAs increase from 11 to 15, the volume increase by 56.2% and surface area increase by 34.0%. The changes in volume, surface area and shape of AABPU due to the mutation provides an overall picture of change in the geometry and structure due to mutation indicating the key role of the inter AAs interaction based on interatomic bonding. In **Figure 3**, we show the details of changes in the volume and shape for mutation Q498R in 4 different orientations with the AAs in the unit marked. It seems obvious that the significant mutation-driven change in all aspects for this AABPU could be one of the reasons leading to the high infectivity of Omicron to be discussed in **Section 4**.

**Figure 3:**
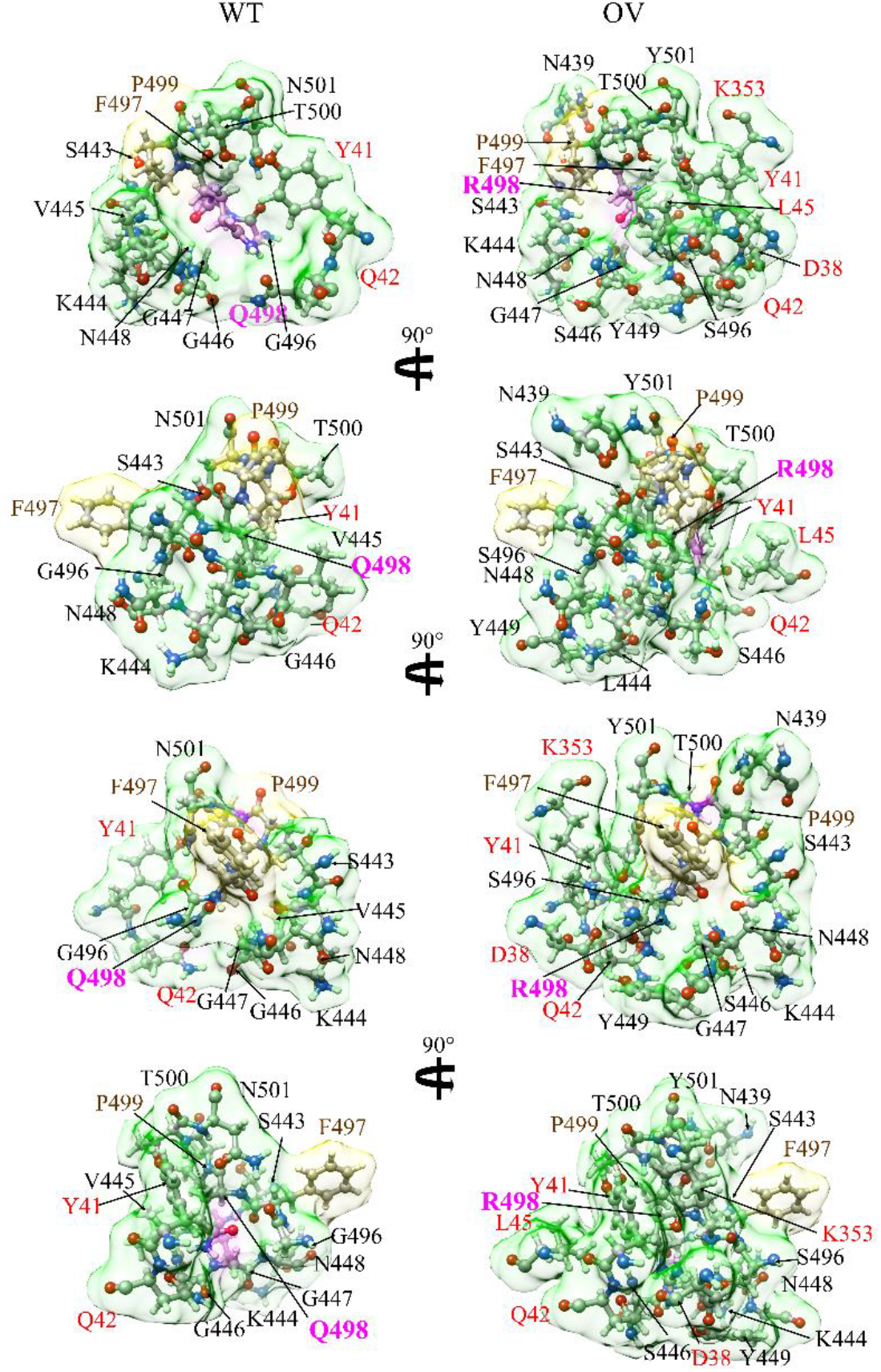
Interaction change due to mutation at site 498 for both WT (left) and OV (right). The surface of site 498 which is marked in magenta color. The surfaces of its NN AAs and NL AAs are shown in yellow and green respectively. Visible NN and NL AAs are marked near to their surface in brown and black respectively. NL AAs from ACE2 are marked in red. We can see considerable changes in NL AAs and their shapes due to mutation.

### 3.2 Electronic structure and bonding

Traditionally, the electronic structure in materials science and condensed matter physics is discussed in the context of the total density of states (TDOS) and the partial density of states (PDOS) of its different components. **Figure S2** shows the TDOS for both WT and the OV from -25 eV to 25 eV. The overall features of the TDOS for WT and OV are very similar since they contain similar atomic components. Both have very low gap between highest occupied molecular orbital (HOMO) and lowest unoccupied molecular orbital (LUMO).

One of the advantages of the OLCAO method is in providing details of the strength of bond between every pair of atoms involved in the system. In **Figure S3 (a)** and **(b)**, we display the bond order (BO) vs. bond length (BL) for every pair of atoms in WT and OV of the RBM-ACE2 model, respectively. The atomic pairs with very short bond length, of 1 Å to 1.1 Å, are N-H, O-H, and C-H bonds. C-O bonds with 1.2 Å to 1.4 Å BL have highest BO, which fluctuates from 0.63 e^-^ to 0.19 e^-^. In the similar range of BL of 1.3 Å to 1.5 Å, the N-C BO ranges from 0.57 e^-^ to 0.12 e^-^. Similarly, from 1.4 Å to 1.7 Å C-C BL can be roughly separated into two groups, one with higher BO from 0.65e^-^ to 0.54e^-^ and the other with lower BO, with the higher BO pair stemming from the double bonds and those with lower BO from the single bonds. There are also notable C-S bonds, with S (Sulphur) from AAs such as Cys, Met, and His. We can see significant amount of HBs in the form of N…H and O…H, with much lower BO and larger BL. There are a few O-Na bonds at BL from 2 Å to 2.3 Å.

While the overall bonding between WT and OV looks similar, there are, however, some differences such as WT has 19 types of different bonds whereas OV has 18 types, with N-Na bonds completely missing in OV. We also notice a significantly stronger C-Na bond between C from I88 and Na ion in the OV case. In the inset for **Figure S3 (a)** and **(b)**, which shows the much smaller BO values at the larger BL from 2.5 Å to 4.5 Å, we notice significant number of C-C, C-H, N-C, H-H, and HBs. Even though these bonds have lower BO, their large number make them significant. The differences between WT in (a) and OV in (b) are present but difficult to be clearly delineate at atomic scale.

### 3.3 Interaction of RBM with ACE2

We now switch our focus to the interface between RBM and ACE2. **Figure 4 (a)** summarizes the interface interaction between the mutated sites of RBM and ACE2. In the WT, 7 out of 10 AAs interact with the ACE2, while in the OV only 6 mutated AAs (S446, R493, S496, R498, Y501, H505) interact with the ACE2. After the mutation in OV A484 loses its bonding with the ACE2. There is a slight decrease in AABP values of S446 with ACE2, indicating a decrease in the interaction. Nevertheless, there are new bonding pairs formed by R493, R498, Y501, and H505, significantly increasing the strength in the RBM-ACE2 interaction. S496, even though interacting with the same components of ACE2, has a slightly increased AABP value, also denoting the increase in the strength of interaction with ACE2. Based on the bonding with ACE2, we contend that these 6 mutated AAs just described can be more critical, when compared to the effect of the remaining mutated AAs, with 5 out of 6 mutated AAs engender a higher AABP value in the bonding with ACE2, indicating an increased binding of RBM to ACE2.

**Figure 4.**
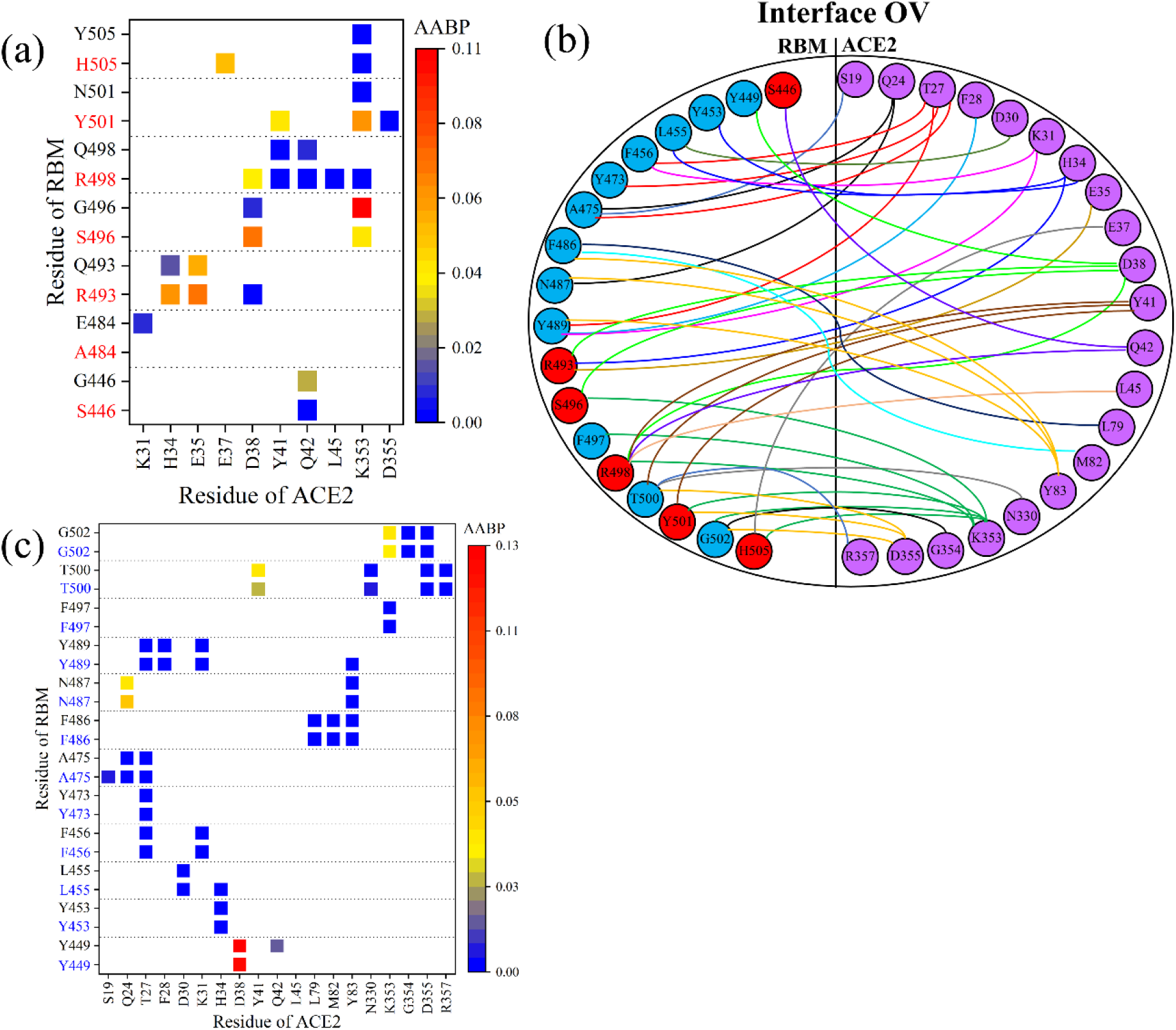
Comparison of RBM-ACE2 interaction in the interface model for WT and OV. (a) AABP map focusing on changes in bonding at the interface of mutated sites for WT (black) and OV (red). (b) interactions at the RBM-ACE2 interface for OV. Interface interacting AAs in RBM are shown in blue and red circles, denoting mutated and unmutated AAs, respectively. Similarly, interacting AAs of ACE2 are shown in purple color. Different colored lines are used just for clarity. (c) AABP map focusing on changes in bonding of interface of unmutated sites for WT (black) and OV (blue).

Only the changes in the bonding of mutated AAs have been discussed so far, but there are also many other unmutated AAs in the RBM that interact with ACE2, which should also be a point of scrutiny. It would thus be of interest to see if there are any changes in the interaction between the unmutated AAs in RBM with ACE2. **Figure 4(b)** shows the complicated topology of the interaction between RBM and ACE2 of the OV in its entirety. This sketch includes the 12 unmutated AAs (blue circles) in addition to 6 mutated AAs (red circles) in the RBM. **Figure 4(c)** shows detailed AABP map for unmutated AAs for both WT and OV. Among the 12 unmutated AAs in OV, it is only the Y449 that has lost its bonding pair with ACE2, while T500 exhibits a reduction in AABP value with Y41 of ACE2. However, the 3 unmutated AAs (L455, A475, and Y489) have increased their bonding with ACE2, so that the changes due to mutation also effects bonding of unmutated AAs, which may also play its part in the increased transmissibility of the OV.

### 3.4 Partial charge of AABPU

Partial charge (PC) is the most important parameter to represent the electrostatic potential around a molecule and is instrumental in predicting the overall long-range intermolecular interaction [48]. From the calculated PC for each atom, we sum them up for all interacting AAs in AABPU for the ten mutations in RBM of OV in **Table 1** and plotted on **Figure 5 (a)** denoted by PC*. For example, the PC* for site N440 is - 1.096 e^-^ obtained from the sum of PC of all interacting AAs N339, N440, L441, S438, D442, and S443. In the OV interface model, the PC* for the mutated K440 is -0.402 e^-^ by summing the PCs of N439, K440, L441, S438, D442, and S443. This is a significant increase toward the positive PC or equivalently a reduction in the negative PC for the mutation N440K. **Figure 5 (a)** shows the comparison of PC* for the 10 sites AABPU before and after mutation. Interestingly, 9 out of 10 sites exhibit an increase in PC* to a more positive charge. The only site that changed from positive PC to negative is Y505H, and even that one only slightly. Ostensibly, the biggest increase in the positive charge is in mutation Q498R from +0.926 e^-^ to 1.889 e^-^, which is more than double. **Figure 5(b)** shows the comparison of PC per AAs (PC^AA^). It shows 10 sites before and after mutation, with 8 out of 10 AAs exhibiting an increased PC in to a more positive charge. Our findings align well with some other indications that PC as a result of mutations may increase [49]. While there have been other studies which have emphasized the dominance of positively charged AAs implicated in the increased electrostatic interactions [49-51], we provide the only quantitative estimate of the effect of the mutations on the PC derived from *ab initio* computation.

**Figure 5.**
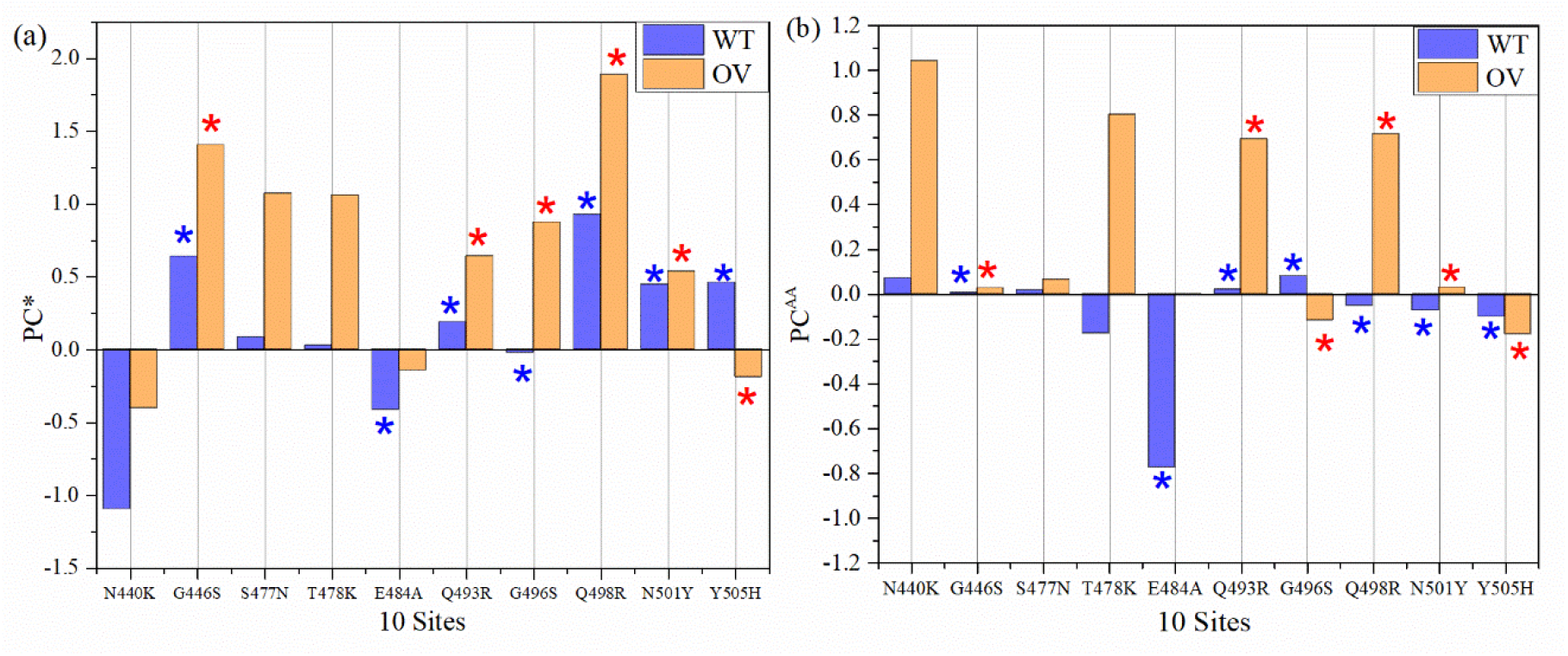
Partial charge (PC) for 10 sites before and after mutation (a) PC per AABPU (PC*) for 10 sites (b) PC per AAs (PC^AA^). 1 out of 10 mutated AABPU (Y505H) flips from positive PC* to negative PC* in relatively small amount. The PC* for mutation Q498R AABPU more than doubled. 2 out of 10 AAs (G496S and Y505H) PC^AA^ value increases in negative direction. The blue and red (*) sign denotes AAs in WT and OV respectively that interacts with ACE2.

It must be emphasized that in **Figure 5 (a)** PC* is for the AABPU, including also the interacting AAs of ACE2. Interestingly, even after interacting with AAs in ACE2 5 out of 6 AABPUs still have a positive PC*, implying that after the interaction with ACE2 the molecular units are more positively charged and may still have further interaction with the negatively charged AAs.

In summing up, all PC* for 10 sites of WT and OV give 1.245 e^-^ and 6.745 e^-^, respectively. The PC* from WT to OV shifts substantially towards positive charge with an impressive increase of 442%. This huge gain in PC* could be one of the major reasons for the rapid infectivity of OV, to be discussed further in **Section 4**. Finally, in **Figure 6**, we correlate the PC* of the 10 mutations with the volume of their AABPU. There is a clearly discernable general trend with mutations tending to make the PC more positive as well as increasing the volume of AABPU, especially and remarkably for the mutation of Q498R.

**Figure 6.**
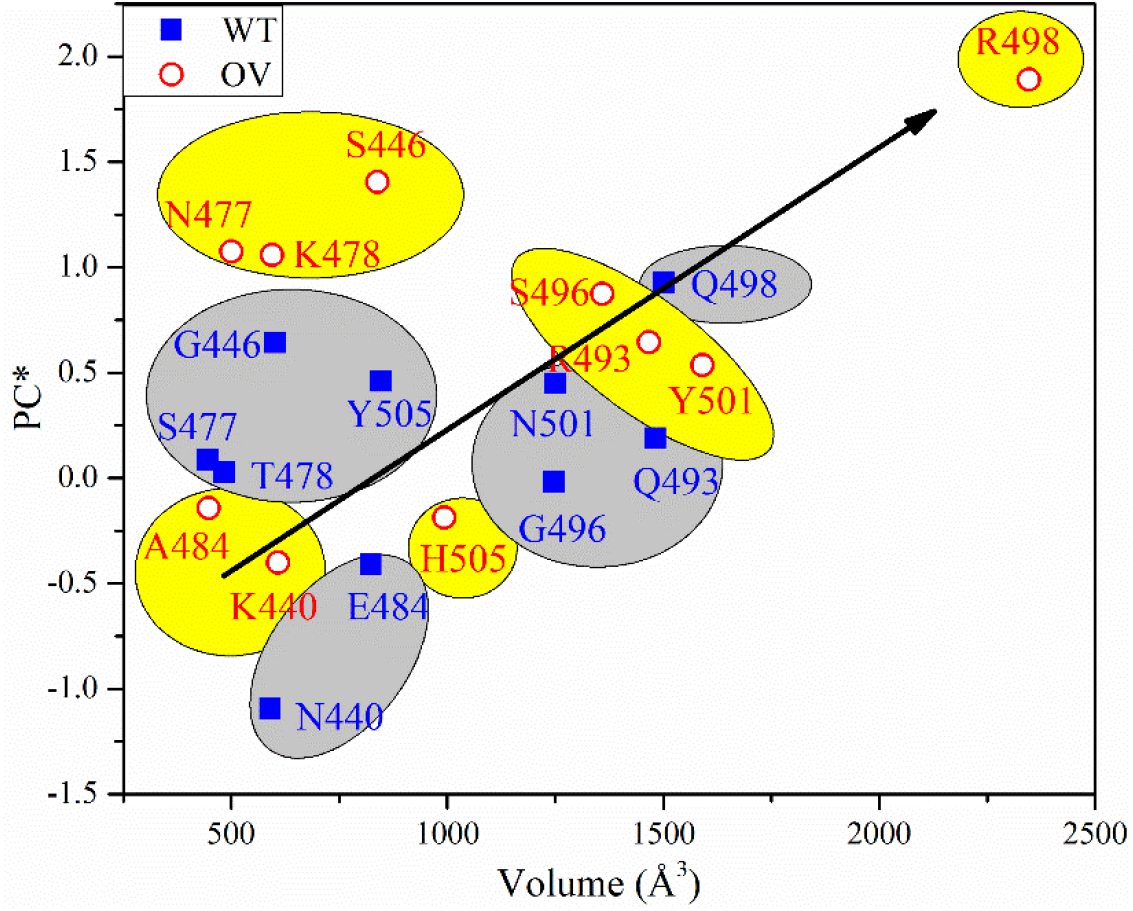
PC* vs volume for 10 sites before and after mutation.

## 4. Source of high infection rate in the Omicron variant

Omicron variant is known for its enhanced infectivity. There are many unanswered questions to this observation, especially at the fundamental level. Based on our detailed analysis of the AABPU data for the 10 mutations of OV, focusing on the changes in their volume and surface areas due to their AAs interactions, we can provide some useful insights. The increase in the total AABP value of the mutated AAs implies the increased interaction within the respective AABPU, whereas a decrease would also denote a decrease of the interaction and consequently an increase in the flexibility in the AABPU. The changes in the number of interacting NL AAs are also critical and will modify the interactions with other AAs, especially those in ACE2, possibly being one of the major reasons of the increased OV infectivity. Among the 10 mutated AAs in OV, 6 interact with ACE2 and 5 of them have an increased AABP value, indicating that the effect of the mutation is to increase the binding of RBM to ACE2. In addition, there is also an increase in AABP values between few unmutated AAs of RBM with ACE2 as an effect of mutation.

Based on our calculated PC values *per* AAs, 8 out of 10 AAs exhibit an increase in the PC^AA^ towards positive charge, confirming the conspicuous dominance of the positively charged AAs as a result of mutations. In addition, our calculations also provided a deeper description of the changes in PC, including the interactions with ACE2. In fact, in 9 out of 10 sites in AABPU, the mutation enforces a positive PC*, which indicates that such AABPU will interact more strongly with negatively charged AAs. Moreover, volume increase of these AABPU follows the increase in their PC*, respectively. The mutations described are mostly limited to the surface of RBM and the overall positive PC* of these sites can thus be related to the high infection rate in the OV, consistent also with studies suggesting development of specific antibodies with mostly negatively charged AAs for better binding [50, 52], as well as the non-specific interactions with predominantly anionic biological membranes [53].

## 5. Conclusion

In summary, we have used the novel concept of AABPU as a basic biomolecular unit in complex proteins to provide detailed information on the effect of 10 mutations in RBM at the interface of RBM-ACE2. For the first time, we also provide accurate values of the volume, surface area, partial charge, and other parameters in AABPU at an atomic level obtained *via* detailed *ab initio* quantum chemical calculations. In particular, the effects of the most important and complicated OV mutation, the Q498R, are clearly described and characterized. This *ab initio* atomic resolution comprehensive investigation of the Omicron variant sheds important light on the fundamental molecular logic behind its enhanced infectivity and paves the way towards a rapid analysis and characterization of the possible next variant(s) of concern in the SARS-CoV-2 virus.

## Supplementary Material

Additional descriptions, figures and Tables are provided in the Supplementary Material.

## Author Contributions

WC and PA conceived the project. WC and PA performed the calculations. PA made most of the figures. WC, and PA, drafted the paper with inputs from BJ and RP. All authors participated in the discussion and interpretation of the results. All authors edited and proofread the final manuscript.

## Institutional Review Board Statement

Not applicable.

## Informed Consent Statement

Not applicable.

## Data Availability Statement

All data are listed in tables or presented in figures in main text or Supplementary Material.

## Acknowledgments

This research used the resources of the National Energy Research Scientific Computing Center supported by DOE under Contract No. DE-AC03-76SF00098 and the Research Computing Support Services (RCSS) of the University of Missouri System. We thank Dr. Richard Gerber, Senior Science Advisor and HPC Department Head for special allocations.

## References

1. Zhong, N., et al., Epidemiology and cause of severe acute respiratory syndrome (SARS) in Guangdong, People’s Republic of China, in February, 2003. The Lancet, 2003. 362(9393): p. 1353–1358, DOI: 10.1016/S0140-6736(03)14630-2.

2. Andrew Rambaut, et al., Preliminary genomic characterisation of an emergent SARS-CoV-2 lineage in the UK defined by a novel set of spike mutations. 2020: SARS-CoV-2 coronavirus nCoV-2019 Genomic Epidemiology.

3. Tegally, H., et al., Emergence and rapid spread of a new severe acute respiratory syndrome-related coronavirus 2 (SARS-CoV-2) lineage with multiple spike mutations in South Africa. MedRxiv, 2020, DOI: 10.1101/2020.12.21.20248640.

4. Singh, J., et al., SARS-CoV-2 variants of concern are emerging in India. Nature medicine, 2021: p. 1–3, DOI: 10.1038/s41591-021-01397-4.

5. Faria, N.R., et al., Genomic characterisation of an emergent SARS-CoV-2 lineage in Manaus: preliminary findings. Virological, 2021. 372: p. 815–821.

6. Ozer, E.A., et al., High prevalence of SARS-CoV-2 B. 1.1. 7 (UK variant) and the novel B. 1.5. 2.5 lineage in Oyo State, Nigeria. medRxiv, 2021, DOI: 10.1101/2021.04.09.21255206.

7. Annavajhala, M.K., et al., A novel SARS-CoV-2 variant of concern, B. 1.526, identified in New York. medRxiv, 2021, DOI: 10.1101/2021.02.23.21252259.

8. Liu, C., et al., Reduced neutralization of SARS-CoV-2 B. 1.617 by vaccine and convalescent serum. Cell, 2021. 184(16): p. 4220-4236. e13, DOI: 10.1016/j.cell.2021.06.020.

9. Kimura, I., et al., SARS-CoV-2 Lambda variant exhibits higher infectivity and immune resistance. bioRxiv, 2021, DOI: 10.1101/2021.07.28.454085.

10. Laiton-Donato, K., et al., Characterization of the emerging B. 1.621 variant of interest of SARS-CoV-2. medRxiv, 2021, DOI: 10.1101/2021.05.08.21256619.

11. Harvey, W.T., et al., SARS-CoV-2 variants, spike mutations and immune escape. Nature Reviews Microbiology, 2021. 19(7): p. 409–424, DOI: 10.1038/s41579-021-00573-0.

12. Omicron Variant: What You Need to Know. 2021 [cited 2022 January 21, 2022]; Available from: https://www.cdc.gov/coronavirus/2019-ncov/variants/omicron-variant.html.

13. Karim, S.S.A. and Q.A. Karim, Omicron SARS-CoV-2 variant: a new chapter in the COVID-19 pandemic. The Lancet, 2021. 398(10317): p. 2126–2128, DOI: 10.1016/S0140-6736(21)02758-6.

14. Science Brief: Omicron (B.1.1.529) Variant. 2021 [cited 2022 January 27]; Available from: https://www.cdc.gov/coronavirus/2019-ncov/science/science-briefs/scientific-brief-omicron-variant.html#print.

15. Classification of Omicron (B.1.1.529): SARS-CoV-2 Variant of Concern. [cited 2022 January 21]; Available from: https://www.who.int/news/item/26-11-2021-classification-of-omicron-(b.1.1.529)-sars-cov-2-variant-of-concern.

16. COVID Data Tracker Weekly Review. [cited 2022 January 21]; Available from: https://www.cdc.gov/coronavirus/2019-ncov/covid-data/covidview/index.html.

17. Torjesen, I., Covid-19: Omicron may be more transmissible than other variants and partly resistant to existing vaccines, scientists fear. 2021, British Medical Journal Publishing Group.

18. Yang, J., et al., A vaccine targeting the RBD of the S protein of SARS-CoV-2 induces protective immunity. Nature, 2020. 586(7830): p. 572–577, DOI: 10.1038/s41586-020-2599-8.

19. Mascola, J.R., B.S. Graham, and A.S. Fauci, SARS-CoV-2 viral variants—tackling a moving target. Jama, 2021. 325(13): p. 1261–1262, DOI: 10.1001/jama.2021.2088.

20. Jiang, S., C. Hillyer, and L. Du, Neutralizing antibodies against SARS-CoV-2 and other human coronaviruses. Trends in immunology, 2020, DOI: 10.1016/j.it.2020.03.007.

21. Huang, Y., et al., Structural and functional properties of SARS-CoV-2 spike protein: potential antivirus drug development for COVID-19. Acta Pharmacologica Sinica, 2020: p. 1–9, DOI: 10.1038/s41401-020-0485-4.

22. Jawad, B., et al., Computational Design of Miniproteins as SARS-CoV-2 Therapeutic Inhibitors. International Journal of Molecular Sciences, 2022. 23(2): p. 838, DOI: 10.3390/ijms23020838.

23. Shang, J., et al., Structural basis of receptor recognition by SARS-CoV-2. Nature, 2020. 581(7807): p. 221–224.

24. Lan, J., et al., Structure of the SARS-CoV-2 spike receptor-binding domain bound to the ACE2 receptor. Nature, 2020. 581(7807): p. 215–220, DOI: 10.1038/s41586-020-2180-5.

25. Yan, R., et al., Structural basis for the recognition of SARS-CoV-2 by full-length human ACE2. Science, 2020. 367(6485): p. 1444–1448, DOI: 10.1126/science.abb2762.

26. Hoffmann, M., et al., The Omicron variant is highly resistant against antibody-mediated neutralization–implications for control of the COVID-19 pandemic. Cell, 2021, DOI: 10.1016/j.cell.2021.12.032.

27. Wilhelm, A., et al., Reduced neutralization of SARS-CoV-2 omicron variant by vaccine sera and monoclonal antibodies. MedRxiv, 2021, DOI: 10.1101/2021.12.07.21267432.

28. Lu, L., et al., Neutralization of SARS-CoV-2 Omicron variant by sera from BNT162b2 or Coronavac vaccine recipients. Clinical Infectious Diseases, 2021, DOI: 10.1093/cid/ciab1041.

29. Collie, S., et al., Effectiveness of BNT162b2 vaccine against omicron variant in South Africa. New England Journal of Medicine, 2021, DOI: 10.1056/NEJMc2119270.

30. Tao, K., et al., The biological and clinical significance of emerging SARS-CoV-2 variants. Nature Reviews Genetics, 2021: p. 1–17, DOI: 10.1038/s41576-021-00408-x.

31. McCallum, M., et al., Molecular basis of immune evasion by the Delta and Kappa SARS-CoV-2 variants. Science, 2021. 374(6575): p. 1621–1626, DOI: 10.1126/science.abl8506.

32. Cai, Y., et al., Structural basis for enhanced infectivity and immune evasion of SARS-CoV-2 variants. Science, 2021. 373(6555): p. 642–648, DOI: 10.1126/science.abi9745.

33. Jawad, B., et al., Key interacting residues between RBD of SARS-CoV-2 and ACE2 receptor: Combination of molecular dynamic simulation and density functional calculation. Journal of Chemical Information and Modeling, 2021, DOI: 10.1021/acs.jcim.1c00560.

34. Jorgensen, W.L., et al., Comparison of simple potential functions for simulating liquid water. J. Chem. Phys., 1983. 79(2): p. 926–935, DOI: 10.1063/1.445869.

35. Pearlman, D.A., et al., AMBER, a package of computer programs for applying molecular mechanics, normal mode analysis, molecular dynamics and free energy calculations to simulate the structural and energetic properties of molecules. Comput. Phys. Commun., 1995. 91(1): p. 1–41, DOI: https://doi.org/10.1016/0010-4655(95)00041-D.

36. Shapovalov, M.V. and R.L. Dunbrack Jr, A smoothed backbone-dependent rotamer library for proteins derived from adaptive kernel density estimates and regressions. Structure, 2011. 19(6): p. 844–858.

37. Pettersen, E.F., et al., UCSF Chimera—a visualization system for exploratory research and analysis. Journal of computational chemistry, 2004. 25(13): p. 1605–1612, DOI: 10.1002/jcc.20084.

38. Local refinement of SARS-CoV-2 S-Beta variant (B.1.351) RBD and Angiotensin-converting enzyme 2 (ACE2) ectodomain. [cited 2021 December 18]; Available from: https://www.rcsb.org/structure/7v80.

39. Package, V.-V.A.i.S.,, Available from: https://www.vasp.at/.

40. Ching, W.-Y. and P. Rulis, Electronic Structure Methods for Complex Materials: The orthogonalized linear combination of atomic orbitals. 2012: Oxford University Press.

41. Adhikari, P. and W.-Y. Ching, Amino acid interacting network in the receptor-binding domain of SARS-CoV-2 spike protein. RSC Advances 2020. 10: p. 39831–39841, DOI: 10.1039/d0ra08222h.

42. Adhikari, P., et al., Intra- and intermolecular atomic-scale interactions in the receptor binding domain of SARS-CoV-2 spike protein: implication for ACE2 receptor binding. Physical Chemistry Chemical Physics, 2020. 22(33): p. 18272–18283, DOI: 10.1039/D0CP03145C.

43. Adhikari, P., et al., First-Principles Simulation of Dielectric Function in Biomolecules. Materials, 2021. 14(19): p. 5774, DOI: 10.3390/ma14195774.

44. Ching, W.-Y., et al., Ultra-Large-Scale Ab Initio Quantum Chemical Computation of Bio-Molecular Systems: The Case of Spike Protein of SARS-CoV-2 Virus.. Computational and Structural Biotechnology Journal 2021. 19: p. 1288–1301, DOI: 10.1016/j.csbj.2021.02.004.

45. Mulliken, R.S., Electronic population analysis on LCAO–MO molecular wave functions. I. J. Chem. Phys., 1955. 23(10): p. 1833–1840.

46. Mulliken, R., Electronic population analysis on LCAO–MO molecular wave functions. II. Overlap populations, bond orders, and covalent bond energies. J. Chem. Phys., 1955. 23(10): p. 1841–1846.

47. Adhikari, P. and W.-Y. Ching, Amino acid interacting network in the receptor-binding domain of SARS-CoV-2 spike protein. RSC Adv., 2020. 10(65): p. 39831–39841, DOI: 10.1039/D0RA08222H.

48. French, R.H., et al., Long range interactions in nanoscale science. Reviews of Modern Physics, 2010. 82(2): p. 1887, DOI: 10.1103/RevModPhys.82.1887.

49. Pawłowski, P.H., Additional positive electric residues in the crucial spike glycoprotein S regions of the new SARS-CoV-2 variants. Infection and Drug Resistance, 2021. 14: p. 5099, DOI: 10.2147/IDR.S342068.

50. Nguyen, H., et al., Electrostatic interactions explain the higher binding affinity of the CR3022 antibody for SARS-CoV-2 than the 4A8 antibody. The Journal of Physical Chemistry B, 2021. 125(27): p. 7368–7379, DOI: 10.1021/acs.jpcb.1c03639.

51. Pawlowski, P.H., Charged amino acids may promote coronavirus SARS-CoV-2 fusion with the host ce. Aims Biophysics, 2021. 8(1): p. 111–120, DOI: 10.3934/biophy.2021008.

52. Neamtu, A., et al., Towards an optimal monoclonal antibody with higher binding affinity to the receptor-binding domain of SARS-CoV-2 spike proteins from different variants. bioRxiv, 2022, DOI: 10.1101/2022.01.04.474958.

53. Khunpetch, P., A. Majee, and R. Podgornik, Curvature effects in charge-regulated lipid bilayers. arXiv preprint, 2022, DOI: 2201.05257.

