## supplementary Materials for "Mutations of Omicron variant at the interface of the receptor domain motif and human angiotensin-converting enzyme-2"

**Content:**

**S1. Methods**

**Figures:**
**Figure S1**: Comparison of (a) Total AABP, (b) NN AABP, (c) NL AABP, (d) AABP from HB, and (e) No. of NL AAs for 10 unmutated (WT) and mutated (OV) AAs. (f) Volume, and (g) Surface for 10 sites or AABPU of RBM-ACE2 interface model for WT and OV.
**Figure S2:** TDOS of RBM-ACE2 interface model for WT and OV.
**Figure S3:** BO vs. BL of RBM-ACE2 interface model for WT and OV.

**Tables:**
**Table S1:** N440 with their bonding for WT interface model.
**Table S2:** K440 with their bonding for OV interface model.
**Table S3:** G446 with their bonding for WT interface model.
**Table S4:** S446 with their bonding for OV interface model.
**Table S5:** S477 with their bonding for WT interface model.
**Table S6:** N447 with their bonding for OV interface model.
**Table S7:** T478 with their bonding for WT interface model.
**Table S8:** K478 with their bonding for OV interface model.
**Table S9:** E484 with their bonding for WT interface model.
**Table S10:** A484 with their bonding for OV interface model.
**Table S11:** Q493 with their bonding for WT interface model.
**Table S12:** R493 with their bonding for OV interface model.
**Table S13:** G496 with their bonding for WT interface model.
**Table S14:** S496 with their bonding for OV interface model.
**Table S15:** Q498 with their bonding for WT interface model.
**Table S16:** R498 with their bonding for OV interface model.
**Table S17:** N501 with their bonding for WT interface model.
**Table S18:** Y501 with their bonding for OV interface model.
**Table S19:** Y505 with their bonding for WT interface model.
**Table S20:** H505 with their bonding for OV interface model.

**References**

### S1. Methods:

#### S1.1 Vienna *ab initio* simulation package (VASP):

The two interface models are fully optimized by using Vienna *ab initio* simulation package (VASP) known for its efficiency in structure optimization.[1] We use the projector augmented wave (PAW) method with Perdew-Burke-Ernzerhof (PBE) exchange correlation functional[2] within the generalized gradient approximation (GGA). The input parameters used in VASP are as follows: energy cut-off 500 eV, electronic convergence of $10^{-4}$ eV, force convergence criteria for ionic steps at $-10^{-2}$ eV/Å and a single k-point sampling. For the optimization, there is complete freedom for ionic position but not for cell volume, and cell shape. All VASP relaxations were carried out at the National Energy Research Scientific Computing (NERSC) facility at the Lawrence Berkeley Laboratory with special allocations and at the Research Computing Support Services (RCSS) of the University of Missouri System. The computational resources used for the structural relaxation are quite substantial because of the high accuracy required in the final structure and the slow convergence for the large complex biomolecular systems.

#### S1.2 Orthogonalized linear combination of atomic orbitals (OLCAO):

In house developed orthogonalized linear combination of atomic orbitals (OLCAO) method [3] is used for the electronic structure and interatomic interactions of the two interface models. Using the OLCAO method we calculate the effective charge ($Q^*$) on each atom and the bond order (BO) values $\rho_{\alpha\beta}$ between any pairs of atoms. They are obtained from the *ab initio* wave functions with atomic basis expansion:

$$Q_\alpha^* = \sum_i \sum_{m,occ} \sum_{j,\beta} C_{i\alpha}^{*m} C_{j\beta}^m S_{i\alpha,j\beta} \tag{1}$$

$$\rho_{\alpha\beta} = \sum_{m,occ} \sum_{i,j} C_{i\alpha}^{*m} C_{j\beta}^m S_{i\alpha,j\beta}. \tag{2}$$

In the above equations, $S_{i\alpha,j\beta}$ are the overlap integrals between the $i^{th}$ orbital in $\alpha^{th}$ atom and the $j^{th}$ orbital in the $\beta^{th}$ atom. $C_{j\beta}^m$ are the eigenvector coefficients of the $m^{th}$occupied molecular orbital level. The partial charge (PC) or ($\Delta Q_\alpha = Q_\alpha^0 \ - Q_\alpha^*$ ) is the deviation of the effective charge $Q_\alpha^*$ from the neutral atomic charge $Q_\alpha^0$ on the same atom $\alpha$. The BO quantifies the strength of the bond between two atoms and usually scales with the bond length (BL). The BL should be more accurately interpreted as the distance of separation of the two atoms since the BO value is influenced by the surrounding atoms. The calculation of PC and BO are based on the Mulliken scheme.[4, 5]

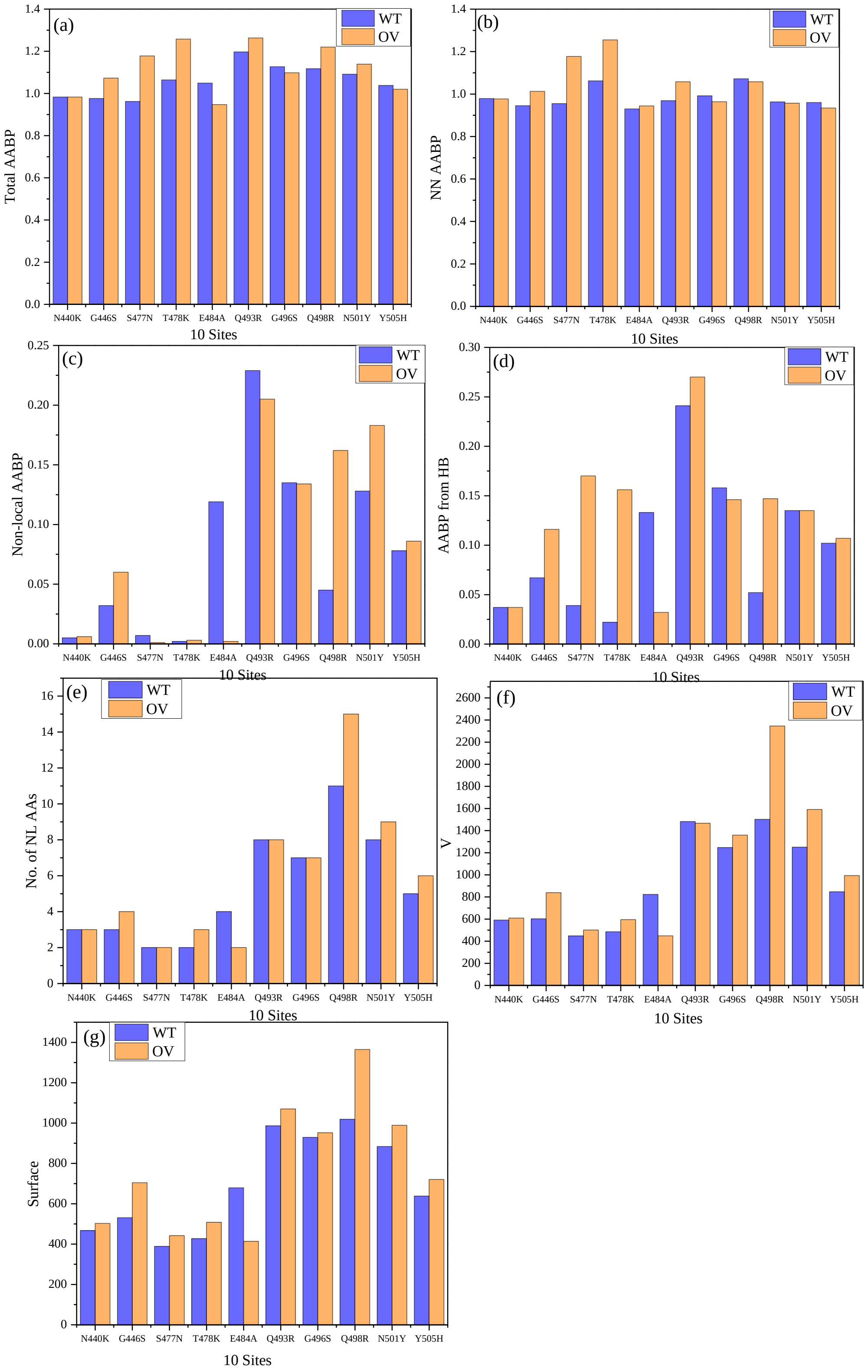


**Figure S1**: Comparison of (a) Total AABP, (b) NN AABP, (c) NL AABP, (d) AABP from HB, and (e) No. of NL AAs for 10 unmutated (WT) and mutated (OV) AAs. (f) Volume, and (g) Surface for 10 sites or AABPU of RBM-ACE2 interface model for WT and OV.

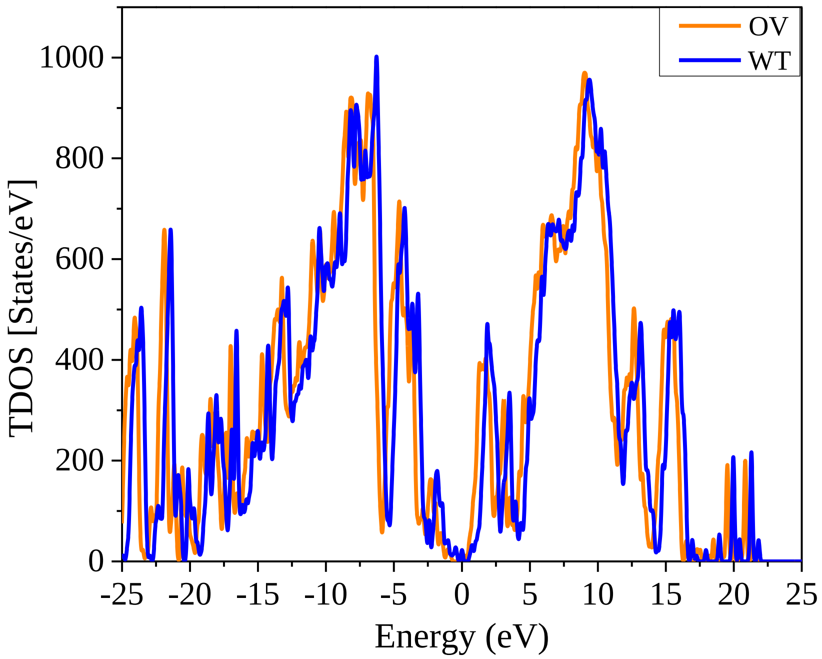


**Figure S2**: TDOS of RBM-ACE2 interface model for WT and OV.

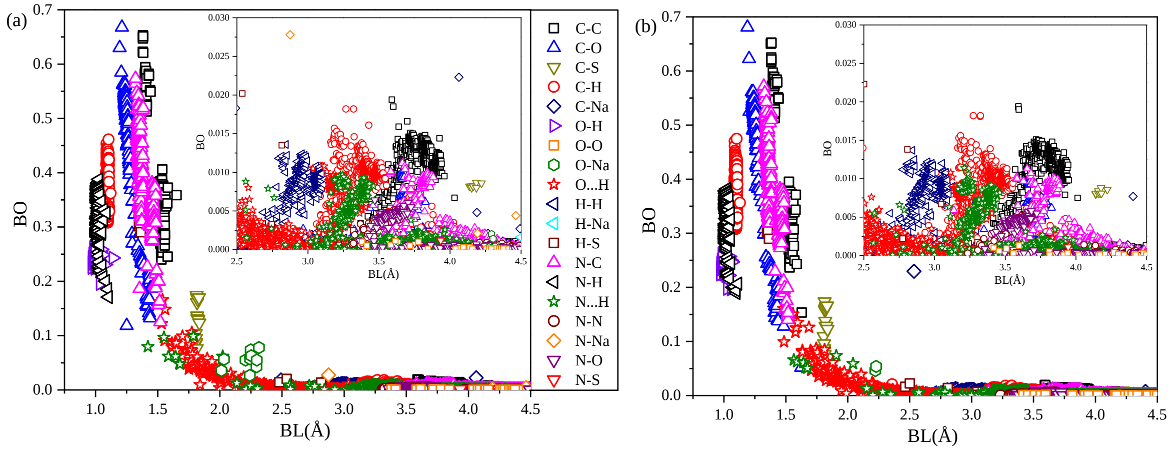


**Figure S3**: BO vs. BL of RBM-ACE2 interface model for **(a)** WT and **(b)** OV.

**Table S1:** N440 with their bonding for WT interface model.

| Bond | BL | BO | AA1 | AA2 |
|---|---|---|---|---|
| N440 | | | | |
| C-C | 3.555 | 0.0047 | N440:C | L441:C |
| C-C | 3.558 | 0.0074 | N440:C | L441:CB |
| C-C | 3.855 | 0.0128 | N440:CA | L441:CA |
| C-O | 3.426 | 0.0008 | N440:C | N439:O |
| C-O | 4.183 | 0.0006 | N440:CB | N439:O |
| H-C | 3.798 | 0.0011 | N440:HB3 | N439:C |
| H-C | 4.071 | 0.0010 | N440:HA | N439:CA |
| H-H | 3.314 | 0.0010 | N440:HA | L441:H |
| N...H | 3.276 | 0.0048 | N440:N | N439:HA |
| N-C | 1.356 | 0.4557 | N440:N | N439:C |
| N-C | 4.191 | 0.0002 | N440:N | N439:CG |
| N-C | 4.235 | 0.0009 | N440:N | L441:CA |
| O...H | 2.427 | 0.0031 | N440:HA | N439:O |
| O...H | 3.193 | 0.0079 | N440:H | N439:O |
| C-C | 3.698 | 0.0122 | N439:C | N440:CB |
| C-C | 3.835 | 0.0133 | N439:CA | N440:CA |
| C-C | 4.324 | 0.0007 | N439:C | N440:CG |
| C-O | 4.118 | 0.0012 | N439:C | N440:OD1 |
| C-O | 3.945 | 0.0010 | L441:CB | N440:O |
| C-O | 3.952 | 0.0002 | L441:CG | N440:O |
| C-O | 3.984 | 0.0007 | L441:C | N440:O |
| H-C | 3.752 | 0.0012 | L441:H | N440:CB |
| H-C | 4.037 | 0.0017 | L441:HA | N440:CA |
| H-C | 4.148 | 0.0002 | L441:HG | N440:CA |
| H-H | 3.578 | 0.0008 | N439:HA | N440:H |
| N...H | 3.111 | 0.0007 | L441:N | N440:HA |
| N...H | 3.926 | 0.0007 | L441:N | N440:HB2 |
| N-C | 4.192 | 0.0009 | N439:N | N440:CA |
| N-C | 1.353 | 0.4199 | L441:N | N440:C |
| N-C | 3.576 | 0.0026 | L441:N | N440:CB |
| O...H | 2.372 | 0.0080 | L441:HA | N440:O |
| O...H | 2.812 | 0.0018 | L441:HD21 | N440:O |
| O...H | 3.194 | 0.0087 | L441:H | N440:O |
| O...H | 4.314 | 0.0001 | L441:HD22 | N440:O |
| O...H | 4.485 | 0.0001 | L441:HD23 | N440:O |
| C-O | 4.060 | 0.0011 | N440:C | S438:O |
| C-O | 4.190 | 0.0005 | N440:CA | S438:O |
| N...H | 4.066 | 0.0001 | N440:N | S438:HA |
| N...H | 4.046 | 0.0014 | N440:N | S443:HG |
| C-C | 4.352 | 0.0002 | S438:C | N440:CA |
| H-C | 3.683 | 0.0010 | D442:H | N440:C |
| H-C | 4.305 | 0.0002 | D442:H | N440:CA |
| N-C | 3.962 | 0.0002 | D442:N | N440:C |

**Table S2:** K440 with their bonding for OV interface model.

| Bond | BL | BO | AA1 | AA2 |
|---|---|---|---|---|
| K440 | | | | |
| C-C | 3.543 | 0.0062 | K440:C | L441:CB |
| C-C | 3.598 | 0.0056 | K440:C | L441:C |
| C-C | 3.870 | 0.0126 | K440:CA | L441:CA |
| C-O | 3.493 | 0.0010 | K440:C | N439:O |
| C-O | 4.209 | 0.0004 | K440:CB | N439:O |
| H-C | 3.799 | 0.0010 | K440:HB3 | N439:C |
| H-C | 4.064 | 0.0005 | K440:HG2 | N439:C |
| H-C | 4.090 | 0.0010 | K440:HA | N439:CA |
| H-C | 4.137 | 0.0003 | K440:HG3 | L441:CA |
| H-C | 4.242 | 0.0003 | K440:HG3 | L441:CB |
| H-C | 4.373 | 0.0002 | K440:HG3 | L441:CD1 |
| H-H | 3.300 | 0.0009 | K440:HA | L441:H |
| N...H | 3.262 | 0.0047 | K440:N | N439:HA |
| N-C | 1.358 | 0.4492 | K440:N | N439:C |
| N-C | 4.186 | 0.0002 | K440:N | N439:CG |
| N-C | 4.251 | 0.0008 | K440:N | L441:CA |
| O...H | 2.465 | 0.0034 | K440:HA | N439:O |
| O...H | 3.150 | 0.0087 | K440:H | N439:O |
| C-C | 3.711 | 0.0121 | N439:C | K440:CB |
| C-C | 3.847 | 0.0131 | N439:CA | K440:CA |
| C-C | 4.422 | 0.0003 | N439:C | K440:CG |
| C-O | 3.895 | 0.0001 | L441:CG | K440:O |
| C-O | 3.923 | 0.0012 | L441:CB | K440:O |
| C-O | 4.052 | 0.0006 | L441:C | K440:O |
| H-C | 3.774 | 0.0012 | L441:H | K440:CB |
| H-C | 4.051 | 0.0004 | L441:HG | K440:CA |
| H-C | 4.059 | 0.0015 | L441:HA | K440:CA |
| H-C | 4.214 | 0.0002 | L441:HG | K440:CB |
| H-C | 4.406 | 0.0003 | L441:HG | K440:CD |
| H-H | 3.462 | 0.0012 | N439:HA | K440:H |
| N...H | 3.109 | 0.0002 | L441:N | K440:HA |
| N...H | 3.884 | 0.0001 | L441:N | K440:HG2 |
| N...H | 3.942 | 0.0007 | L441:N | K440:HB2 |
| N-C | 1.354 | 0.4248 | L441:N | K440:C |
| N-C | 3.592 | 0.0030 | L441:N | K440:CB |
| N-C | 4.190 | 0.0010 | N439:N | K440:CA |
| O...H | 2.406 | 0.0069 | L441:HA | K440:O |
| O...H | 2.784 | 0.0019 | L441:HD21 | K440:O |
| O...H | 3.196 | 0.0089 | L441:H | K440:O |
| O...H | 4.243 | 0.0001 | L441:HD22 | K440:O |
| O...H | 4.475 | 0.0001 | L441:HD23 | K440:O |
| C-O | 3.928 | 0.0016 | K440:C | S438:O |
| C-O | 4.059 | 0.0006 | K440:CA | S438:O |
| N...H | 4.053 | 0.0014 | K440:N | S443:HG |
| N...H | 4.134 | 0.0001 | K440:N | S438:HA |
| C-C | 4.328 | 0.0003 | S438:C | K440:CA |
| H-C | 3.764 | 0.0011 | D442:H | K440:C |
| H-C | 4.414 | 0.0002 | D442:H | K440:CA |
| N-C | 4.045 | 0.0003 | D442:N | K440:C |

**Table S3:** G446 with their bonding for WT interface model.

| G446 | | | | |
|---|---|---|---|---|
| C-C | 3.457 | 0.0018 | G446:C | G447:C |
| C-C | 3.837 | 0.0135 | G446:CA | G447:CA |
| C-O | 3.762 | 0.0009 | G446:C | V445:O |
| C-O | 3.588 | 0.0002 | G446:C | G447:O |
| H-C | 3.279 | 0.0080 | G446:HA2 | V445:C |
| H-C | 3.560 | 0.0013 | G446:H | V445:CB |
| H-C | 4.086 | 0.0015 | G446:HA3 | V445:CA |
| H-C | 4.405 | 0.0001 | G446:H | G447:CA |
| H-H | 3.163 | 0.0004 | G446:HA3 | G447:H |
| H-H | 3.521 | 0.0006 | G446:HA2 | G447:H |
| N...H | 3.916 | 0.0006 | G446:N | V445:HB |
| N...H | 4.471 | 0.0002 | G446:N | V445:HG12 |
| N-C | 1.361 | 0.3932 | G446:N | V445:C |
| N-C | 3.463 | 0.0012 | G446:N | V445:CB |
| N-C | 4.248 | 0.0006 | G446:N | G447:CA |
| O...H | 2.447 | 0.0075 | G446:HA3 | V445:O |
| O...H | 3.175 | 0.0078 | G446:H | V445:O |
| O...H | 3.750 | 0.0001 | G446:HA2 | V445:O |
| C-C | 3.469 | 0.0017 | V445:C | G446:C |
| C-C | 3.837 | 0.0137 | V445:CA | G446:CA |
| C-O | 3.771 | 0.0004 | G447:C | G446:O |
| H-C | 3.945 | 0.0012 | V445:HA | G446:CA |
| H-C | 3.211 | 0.0075 | G447:HA2 | G446:C |
| H-C | 4.033 | 0.0018 | G447:HA3 | G446:CA |
| N...H | 3.668 | 0.0012 | V445:N | G446:H |
| N...H | 3.261 | 0.0030 | G447:N | G446:HA2 |
| N-C | 1.354 | 0.4523 | G447:N | G446:C |
| N-N | 3.501 | 0.0043 | V445:N | G446:N |
| O...H | 2.372 | 0.0095 | G447:HA3 | G446:O |
| O...H | 3.194 | 0.0079 | G447:H | G446:O |
| O...H | 3.663 | 0.0005 | G447:HA2 | G446:O |
| C-O | 4.105 | 0.0008 | G446:C | K444:O |
| C-O | 4.196 | 0.0005 | G446:CA | K444:O |
| H-C | 3.909 | 0.0001 | G446:H | K444:C |
| O...H | 3.314 | 0.0003 | G446:H | K444:O |
| N-C | 3.766 | 0.0007 | G446:N | K444:C |
| H-C | 4.450 | 0.0001 | G446:H | Q498:CD |
| C-O | 4.019 | 0.0009 | Q42:CD | G446:O |
| H-C | 4.392 | 0.0002 | Q42:HE22 | G446:C |
| H-H | 4.317 | 0.0001 | Q42:HE21 | G446:HA3 |
| O...H | 1.997 | 0.0279 | Q42:HE21 | G446:O |
| O...H | 3.362 | 0.0002 | Q42:HE22 | G446:O |
| H-C | 4.061 | 0.0001 | Q498:HG3 | G446:CA |

**Table S4:** S446 with their bonding for OV interface model.

| S446 | | | | |
|---|---|---|---|---|
| C-C | 3.492 | 0.0021 | S446:C | G447:C |
| C-C | 3.844 | 0.0133 | S446:CA | G447:CA |
| C-O | 3.631 | 0.0005 | S446:C | G447:O |
| C-O | 3.906 | 0.0012 | S446:C | V445:O |
| H-C | 3.310 | 0.0081 | S446:HA | V445:C |
| H-C | 3.438 | 0.0014 | S446:H | V445:CB |
| H-C | 3.941 | 0.0022 | S446:HG | V445:CA |
| H-C | 4.413 | 0.0001 | S446:H | G447:CA |
| H-H | 3.316 | 0.0003 | S446:HB2 | G447:H |
| H-H | 3.498 | 0.0007 | S446:HA | G447:H |
| N...H | 3.861 | 0.0006 | S446:N | V445:HB |
| N...H | 4.418 | 0.0002 | S446:N | V445:HG12 |
| N-C | 1.359 | 0.4278 | S446:N | V445:C |
| N-C | 3.432 | 0.0007 | S446:N | V445:CB |
| N-C | 4.201 | 0.0010 | S446:N | G447:CA |
| O...H | 2.081 | 0.0299 | S446:HG | V445:O |
| O...H | 3.165 | 0.0084 | S446:H | V445:O |
| O...H | 3.957 | 0.0009 | S446:HB3 | V445:O |
| C-C | 3.498 | 0.0008 | V445:C | S446:C |
| C-C | 3.875 | 0.0132 | V445:CA | S446:CA |
| C-O | 3.860 | 0.0008 | G447:C | S446:O |
| H-C | 3.209 | 0.0074 | G447:HA2 | S446:C |
| H-C | 3.558 | 0.0005 | G447:H | S446:CB |
| H-C | 3.941 | 0.0012 | V445:HA | S446:CA |
| H-C | 4.072 | 0.0015 | G447:HA3 | S446:CA |
| H-C | 4.113 | 0.0001 | V445:HG13 | S446:CB |
| H-H | 4.316 | 0.0001 | V445:HG12 | S446:H |
| N...H | 3.250 | 0.0034 | G447:N | S446:HA |
| N...H | 3.663 | 0.0014 | V445:N | S446:H |
| N...H | 3.823 | 0.0004 | G447:N | S446:HB3 |
| N-C | 1.354 | 0.4615 | G447:N | S446:C |
| N-N | 3.505 | 0.0043 | V445:N | S446:N |
| O...H | 2.409 | 0.0083 | G447:HA3 | S446:O |
| O...H | 3.200 | 0.0080 | G447:H | S446:O |
| O...H | 3.650 | 0.0004 | G447:HA2 | S446:O |
| C-C | 4.080 | 0.0010 | S446:C | R498:CZ |
| C-O | 4.075 | 0.0010 | S446:C | K444:O |
| C-O | 4.248 | 0.0005 | S446:CA | K444:O |
| H-C | 3.967 | 0.0002 | S446:H | K444:C |
| O...H | 3.412 | 0.0003 | S446:H | K444:O |
| N-C | 3.795 | 0.0007 | S446:N | K444:C |
| C-O | 3.628 | 0.0007 | R498:CZ | S446:O |
| H-C | 3.986 | 0.0016 | R498:HH21 | S446:CA |
| H-C | 4.024 | 0.0001 | Y449:HE2 | S446:C |
| H-C | 4.309 | 0.0001 | R498:HG3 | S446:CA |
| H-C | 4.497 | 0.0002 | R498:HH22 | S446:C |
| O...H | 1.867 | 0.0525 | R498:HH21 | S446:O |
| O...H | 3.029 | 0.0002 | Q42:HE21 | S446:O |
| O...H | 3.537 | 0.0008 | R498:HH22 | S446:O |
| N-C | 3.695 | 0.0004 | R498:NH2 | S446:C |

**Table S5:** S477 with their bonding for WT interface model.

| S477 | | | | |
|---|---|---|---|---|
| C-C | 3.653 | 0.0124 | S477:C | T478:CB |
| C-C | 3.856 | 0.0134 | S477:CA | T478:CA |
| C-O | 3.664 | 0.0004 | S477:C | G476:O |
| C-O | 4.124 | 0.0007 | S477:CB | G476:O |
| C-O | 4.059 | 0.0002 | S477:C | T478:O |
| C-O | 4.069 | 0.0004 | S477:C | T478:OG1 |
| H-C | 3.690 | 0.0011 | S477:HB3 | G476:C |
| H-C | 4.038 | 0.0014 | S477:HA | G476:CA |
| H-C | 4.478 | 0.0002 | S477:H | T478:CA |
| H-H | 3.339 | 0.0007 | S477:HA | T478:H |
| N...H | 3.848 | 0.0010 | S477:N | G476:H |
| N-C | 1.364 | 0.4301 | S477:N | G476:C |
| N-C | 4.219 | 0.0009 | S477:N | T478:CA |
| O...H | 2.424 | 0.0058 | S477:HA | G476:O |
| O...H | 3.186 | 0.0078 | S477:H | G476:O |
| C-C | 3.462 | 0.0018 | G476:C | S477:C |
| C-C | 3.621 | 0.0109 | G476:C | S477:CB |
| C-C | 3.825 | 0.0126 | G476:CA | S477:CA |
| C-O | 4.065 | 0.0004 | G476:C | S477:OG |
| C-O | 3.674 | 0.0010 | T478:C | S477:O |
| C-O | 4.234 | 0.0004 | T478:CB | S477:O |
| C-O | 4.493 | 0.0001 | T478:CG2 | S477:O |
| H-C | 3.997 | 0.0010 | G476:HA3 | S477:CA |
| H-C | 3.729 | 0.0014 | T478:H | S477:CB |
| H-C | 4.107 | 0.0012 | T478:HA | S477:CA |
| H-C | 4.155 | 0.0001 | T478:HG21 | S477:CA |
| H-C | 4.215 | 0.0002 | T478:HG23 | S477:C |
| H-C | 4.219 | 0.0001 | T478:HG1 | S477:C |
| H-H | 3.009 | 0.0006 | G476:HA2 | S477:H |
| N...H | 3.883 | 0.0011 | G476:N | S477:H |
| N...H | 3.140 | 0.0008 | T478:N | S477:HA |
| N...H | 3.808 | 0.0011 | T478:N | S477:HB2 |
| N-C | 1.347 | 0.4227 | T478:N | S477:C |
| N-C | 3.533 | 0.0019 | T478:N | S477:CB |
| N-N | 3.625 | 0.0043 | G476:N | S477:N |
| O...H | 2.538 | 0.0063 | T478:HA | S477:O |
| O...H | 3.192 | 0.0084 | T478:H | S477:O |
| N...H | 4.432 | 0.0001 | S477:N | Q474:HG2 |
| N-N | 4.455 | 0.0003 | Q474:NE2 | S477:N |
| O...H | 2.561 | 0.0063 | P479:HD3 | S477:O |
| O...H | 4.173 | 0.0001 | P479:HD2 | S477:O |
| O...H | 4.341 | 0.0001 | P479:HG3 | S477:O |
| N-C | 4.036 | 0.0003 | P479:N | S477:C |

**Table S6:** N447 with their bonding for OV interface model.

| N477 | | | | |
|---|---|---|---|---|
| C-C | 3.530 | 0.0042 | N477:C | K478:CB |
| C-C | 3.555 | 0.0087 | N477:C | K478:C |
| C-C | 3.792 | 0.0010 | N477:C | K478:CG |
| C-C | 3.842 | 0.0126 | N477:CA | K478:CA |
| C-C | 4.027 | 0.0010 | N477:C | K478:CE |
| C-O | 3.886 | 0.0002 | N477:C | G476:O |
| C-O | 3.899 | 0.0012 | N477:CB | G476:O |
| C-O | 4.036 | 0.0005 | N477:C | K478:O |
| H-C | 3.388 | 0.0004 | N477:HB3 | G476:C |
| H-C | 3.974 | 0.0021 | N477:HA | G476:CA |
| H-C | 4.084 | 0.0001 | N477:HA | K478:CA |
| N...H | 3.828 | 0.0009 | N477:N | G476:H |
| N-C | 1.371 | 0.4295 | N477:N | G476:C |
| N-C | 4.328 | 0.0005 | N477:N | K478:CA |
| N-N | 4.167 | 0.0001 | N477:ND2 | K478:NZ |
| O...H | 2.287 | 0.0104 | N477:HA | G476:O |
| O...H | 3.196 | 0.0075 | N477:H | G476:O |
| O...H | 3.717 | 0.0001 | N477:HB3 | G476:O |
| C-C | 3.506 | 0.0043 | G476:C | N477:CB |
| C-C | 3.576 | 0.0074 | G476:C | N477:C |
| C-C | 3.819 | 0.0123 | G476:CA | N477:CA |
| C-C | 4.305 | 0.0002 | G476:C | N477:CG |
| C-O | 3.571 | 0.0014 | K478:CE | N477:O |
| C-O | 3.794 | 0.0006 | K478:CB | N477:O |
| C-O | 4.138 | 0.0004 | K478:C | N477:O |
| C-O | 4.252 | 0.0009 | G476:C | N477:OD1 |
| H-C | 3.598 | 0.0028 | K478:HZ1 | N477:CA |
| H-C | 3.788 | 0.0003 | K478:HG3 | N477:C |
| H-C | 3.838 | 0.0002 | K478:HB2 | N477:C |
| H-C | 3.994 | 0.0011 | G476:HA3 | N477:CA |
| H-C | 4.036 | 0.0010 | K478:HA | N477:CA |
| H-C | 4.045 | 0.0006 | K478:HZ3 | N477:C |
| H-C | 4.067 | 0.0011 | K478:H | N477:CB |
| H-C | 4.126 | 0.0004 | K478:HZ2 | N477:C |
| H-C | 4.269 | 0.0002 | K478:HE2 | N477:C |
| H-C | 4.370 | 0.0003 | K478:HD3 | N477:C |
| H-H | 3.086 | 0.0008 | G476:HA2 | N477:H |
| N...H | 3.867 | 0.0011 | G476:N | N477:H |
| N...H | 4.057 | 0.0005 | K478:N | N477:HB2 |
| N-C | 1.343 | 0.4977 | K478:N | N477:C |
| N-C | 3.543 | 0.0003 | K478:NZ | N477:C |
| N-C | 3.755 | 0.0062 | K478:N | N477:CB |
| N-C | 4.239 | 0.0004 | K478:N | N477:CG |
| N-N | 3.613 | 0.0041 | G476:N | N477:N |
| O...H | 1.596 | 0.1350 | K478:HZ1 | N477:O |
| O...H | 2.458 | 0.0031 | K478:HA | N477:O |
| O...H | 2.723 | 0.0016 | K478:HE2 | N477:OD1 |
| O...H | 2.846 | 0.0015 | K478:HD2 | N477:OD1 |
| O...H | 3.199 | 0.0079 | K478:H | N477:O |
| O...H | 3.301 | 0.0001 | K478:HZ2 | N477:OD1 |
| O...H | 4.082 | 0.0001 | K478:HD3 | N477:OD1 |
| O...H | 4.302 | 0.0002 | K478:HZ3 | N477:OD1 |
| O...H | 4.348 | 0.0001 | K478:HE3 | N477:OD1 |
| O...H | 4.486 | 0.0001 | K478:HD3 | N477:O |
| H-C | 3.945 | 0.0002 | P479:HD3 | N477:C |
| H-C | 4.341 | 0.0001 | Q474:HE22 | N477:C |
| N-C | 4.431 | 0.0001 | P479:N | N477:C |
| N-C | 4.464 | 0.0001 | Q474:NE2 | N477:CA |

**Table S7:** T478 with their bonding for WT interface model.

| T478 | | | | |
|---|---|---|---|---|
| C-C | 3.638 | 0.0111 | T478:C | P479:CG |
| C-C | 3.662 | 0.0134 | T478:C | P479:CB |
| C-C | 3.857 | 0.0125 | T478:CA | P479:CA |
| C-C | 4.215 | 0.0014 | T478:CB | P479:CD |
| C-C | 4.405 | 0.0004 | T478:CA | P479:C |
| C-C | 4.421 | 0.0008 | T478:CA | P479:CG |
| C-O | 3.674 | 0.0010 | T478:C | S477:O |
| C-O | 4.234 | 0.0004 | T478:CB | S477:O |
| C-O | 4.493 | 0.0001 | T478:CG2 | S477:O |
| H-C | 3.729 | 0.0014 | T478:H | S477:CB |
| H-C | 4.107 | 0.0012 | T478:HA | S477:CA |
| H-C | 4.155 | 0.0001 | T478:HG21 | S477:CA |
| H-C | 4.215 | 0.0002 | T478:HG23 | S477:C |
| H-C | 4.219 | 0.0001 | T478:HG1 | S477:C |
| H-C | 4.044 | 0.0013 | T478:HA | P479:CG |
| H-C | 4.075 | 0.0005 | T478:HA | P479:CA |
| H-C | 4.090 | 0.0001 | T478:HB | P479:CD |
| H-H | 3.840 | 0.0001 | T478:HB | P479:HD2 |
| N...H | 3.140 | 0.0008 | T478:N | S477:HA |
| N...H | 3.808 | 0.0011 | T478:N | S477:HB2 |
| N-C | 1.347 | 0.4227 | T478:N | S477:C |
| N-C | 3.533 | 0.0019 | T478:N | S477:CB |
| N-C | 3.932 | 0.0020 | T478:N | P479:CD |
| N-N | 3.464 | 0.0017 | T478:N | P479:N |
| O...H | 2.538 | 0.0063 | T478:HA | S477:O |
| O...H | 3.192 | 0.0084 | T478:H | S477:O |
| O...H | 3.685 | 0.0001 | T478:HB | P479:O |
| O...H | 4.339 | 0.0001 | T478:HG1 | P479:O |
| C-C | 3.653 | 0.0124 | S477:C | T478:CB |
| C-C | 3.856 | 0.0134 | S477:CA | T478:CA |
| C-O | 4.059 | 0.0002 | S477:C | T478:O |
| C-O | 4.069 | 0.0004 | S477:C | T478:OG1 |
| C-O | 3.666 | 0.0091 | P479:CD | T478:O |
| C-O | 4.233 | 0.0010 | P479:CB | T478:O |
| H-C | 4.478 | 0.0002 | S477:H | T478:CA |
| H-C | 3.914 | 0.0005 | P479:HG3 | T478:C |
| H-C | 4.163 | 0.0002 | P479:HD3 | T478:CB |
| H-C | 4.171 | 0.0003 | P479:HD2 | T478:CB |
| H-H | 3.339 | 0.0007 | S477:HA | T478:H |
| N...H | 3.425 | 0.0001 | P479:N | T478:HB |
| N...H | 3.766 | 0.0023 | P479:N | T478:HG1 |
| N-C | 4.219 | 0.0009 | S477:N | T478:CA |
| N-C | 1.347 | 0.5251 | P479:N | T478:C |
| N-C | 3.569 | 0.0025 | P479:N | T478:CB |
| O...H | 2.769 | 0.0005 | P479:HA | T478:O |
| O...H | 3.961 | 0.0005 | P479:HD3 | T478:O |
| O...H | 4.168 | 0.0002 | P479:HD2 | T478:O |
| C-O | 4.091 | 0.0001 | T478:C | Q474:OE1 |
| H-C | 4.490 | 0.0001 | Q474:HG3 | T478:CA |
| O...H | 2.583 | 0.0007 | Q474:HG3 | T478:O |
| O...H | 4.065 | 0.0002 | Q474:HG2 | T478:O |
| O...H | 4.469 | 0.0001 | Q474:HB2 | T478:O |
| O...H | 3.508 | 0.0002 | C480:H | T478:O |
| O...H | 3.996 | 0.0002 | C480:HA | T478:O |
| N-C | 4.023 | 0.0006 | C480:N | T478:C |

**Table S8:** K478 with their bonding for OV interface model.

| K478 | | | | |
|---|---|---|---|---|
| C-C | 3.619 | 0.0104 | K478:C | P479:CG |
| C-C | 3.679 | 0.0131 | K478:C | P479:CB |
| C-C | 3.858 | 0.0124 | K478:CA | P479:CA |
| C-C | 3.973 | 0.0014 | K478:CB | P479:CD |
| C-C | 4.401 | 0.0007 | K478:CA | P479:CG |
| C-C | 4.405 | 0.0002 | K478:CA | P479:C |
| C-O | 3.571 | 0.0014 | K478:CE | N477:O |
| C-O | 3.794 | 0.0006 | K478:CB | N477:O |
| C-O | 4.138 | 0.0004 | K478:C | N477:O |
| H-C | 3.598 | 0.0028 | K478:HZ1 | N477:CA |
| H-C | 3.788 | 0.0003 | K478:HG3 | N477:C |
| H-C | 3.838 | 0.0002 | K478:HB2 | N477:C |
| H-C | 4.014 | 0.0013 | K478:HA | P479:CG |
| H-C | 4.036 | 0.0010 | K478:HA | N477:CA |
| H-C | 4.045 | 0.0006 | K478:HZ3 | N477:C |
| H-C | 4.067 | 0.0011 | K478:H | N477:CB |
| H-C | 4.108 | 0.0006 | K478:HA | P479:CA |
| H-C | 4.126 | 0.0004 | K478:HZ2 | N477:C |
| H-C | 4.269 | 0.0002 | K478:HE2 | N477:C |
| H-C | 4.370 | 0.0003 | K478:HD3 | N477:C |
| H-H | 4.030 | 0.0001 | K478:HG3 | P479:HD3 |
| H-H | 4.224 | 0.0001 | K478:HA | P479:HG3 |
| N...H | 3.616 | 0.0007 | K478:N | P479:HD3 |
| N...H | 4.057 | 0.0005 | K478:N | N477:HB2 |
| N-C | 1.343 | 0.4977 | K478:N | N477:C |
| N-C | 3.543 | 0.0003 | K478:NZ | N477:C |
| N-C | 3.755 | 0.0062 | K478:N | N477:CB |
| N-C | 4.092 | 0.0017 | K478:N | P479:CD |
| N-C | 4.239 | 0.0004 | K478:N | N477:CG |
| N-N | 3.492 | 0.0027 | K478:N | P479:N |
| O...H | 1.596 | 0.1350 | K478:HZ1 | N477:O |
| O...H | 2.458 | 0.0031 | K478:HA | N477:O |
| O...H | 2.723 | 0.0016 | K478:HE2 | N477:OD1 |
| O...H | 2.846 | 0.0015 | K478:HD2 | N477:OD1 |
| O...H | 3.199 | 0.0079 | K478:H | N477:O |
| O...H | 3.301 | 0.0001 | K478:HZ2 | N477:OD1 |
| O...H | 3.341 | 0.0001 | K478:HB3 | P479:O |
| O...H | 4.082 | 0.0001 | K478:HD3 | N477:OD1 |
| O...H | 4.302 | 0.0002 | K478:HZ3 | N477:OD1 |
| O...H | 4.348 | 0.0001 | K478:HE3 | N477:OD1 |
| O...H | 4.486 | 0.0001 | K478:HD3 | N477:O |
| C-C | 3.530 | 0.0042 | N477:C | K478:CB |
| C-C | 3.555 | 0.0087 | N477:C | K478:C |
| C-C | 3.792 | 0.0010 | N477:C | K478:CG |
| C-C | 3.842 | 0.0126 | N477:CA | K478:CA |
| C-C | 4.027 | 0.0010 | N477:C | K478:CE |
| C-O | 3.671 | 0.0092 | P479:CD | K478:O |
| C-O | 4.036 | 0.0005 | N477:C | K478:O |
| C-O | 4.261 | 0.0008 | P479:CB | K478:O |
| H-C | 3.846 | 0.0006 | P479:HG3 | K478:C |
| H-C | 3.897 | 0.0004 | P479:HD3 | K478:CB |
| H-C | 3.915 | 0.0003 | P479:HD2 | K478:CB |
| H-C | 4.084 | 0.0001 | N477:HA | K478:CA |
| H-C | 4.198 | 0.0003 | P479:HA | K478:CA |
| N...H | 3.678 | 0.0015 | P479:N | K478:H |
| N...H | 3.922 | 0.0003 | P479:N | K478:HB2 |
| N-C | 1.351 | 0.5016 | P479:N | K478:C |
| N-C | 4.328 | 0.0005 | N477:N | K478:CA |
| N-N | 4.167 | 0.0001 | N477:ND2 | K478:NZ |
| O...H | 2.663 | 0.0007 | P479:HA | K478:O |
| O...H | 3.965 | 0.0005 | P479:HD3 | K478:O |
| O...H | 4.202 | 0.0001 | P479:HD2 | K478:O |
| N-O | 3.912 | 0.0001 | K478:N | G476:O |
| C-C | 4.318 | 0.0002 | Q474:CG | K478:C |
| C-O | 4.474 | 0.0004 | Q474:CB | K478:O |
| H-C | 4.174 | 0.0001 | C480:H | K478:C |
| N-C | 4.091 | 0.0005 | C480:N | K478:C |
| O...H | 2.458 | 0.0016 | Q474:HG3 | K478:O |
| O...H | 3.687 | 0.0001 | C480:H | K478:O |
| O...H | 3.906 | 0.0002 | Q474:HG2 | K478:O |
| O...H | 4.353 | 0.0001 | C480:HA | K478:O |
| O...H | 4.378 | 0.0001 | Q474:HB2 | K478:O |

**Table S9:** E484 with their bonding for WT interface model.

| E484 | | | | |
|---|---|---|---|---|
| C-C | 3.721 | 0.0139 | E484:C | G485:C |
| C-C | 3.796 | 0.0143 | E484:CA | G485:CA |
| C-O | 3.512 | 0.0013 | E484:C | V483:O |
| C-O | 4.263 | 0.0003 | E484:CB | V483:O |
| H-C | 3.684 | 0.0005 | E484:H | V483:CG2 |
| H-C | 4.010 | 0.0004 | E484:H | V483:CG1 |
| H-C | 4.044 | 0.0005 | E484:HB2 | V483:C |
| H-C | 4.123 | 0.0009 | E484:HA | V483:CA |
| H-C | 3.977 | 0.0010 | E484:HA | G485:CA |
| N...H | 4.081 | 0.0006 | E484:N | V483:H |
| N...H | 3.901 | 0.0011 | E484:N | G485:H |
| N-C | 1.365 | 0.4205 | E484:N | V483:C |
| N-C | 3.718 | 0.0002 | E484:N | V483:CG2 |
| N-C | 4.440 | 0.0002 | E484:N | V483:CG1 |
| N-N | 3.633 | 0.0052 | E484:N | G485:N |
| O...H | 2.503 | 0.0036 | E484:HA | V483:O |
| O...H | 3.170 | 0.0082 | E484:H | V483:O |
| C-C | 3.762 | 0.0122 | V483:C | E484:CB |
| C-C | 3.836 | 0.0124 | V483:CA | E484:CA |
| C-C | 4.363 | 0.0002 | V483:CB | E484:CA |
| C-O | 3.563 | 0.0001 | V483:C | E484:O |
| C-O | 4.256 | 0.0004 | G485:C | E484:O |
| H-C | 3.908 | 0.0004 | V483:HB | E484:CA |
| H-C | 4.118 | 0.0006 | V483:HA | E484:CA |
| H-C | 4.139 | 0.0005 | G485:HA2 | E484:CA |
| H-C | 4.191 | 0.0004 | G485:HA3 | E484:CA |
| H-C | 4.453 | 0.0002 | G485:HA2 | E484:CB |
| H-H | 3.428 | 0.0001 | V483:HG21 | E484:H |
| H-H | 4.017 | 0.0001 | V483:HG13 | E484:H |
| N...H | 3.956 | 0.0009 | V483:N | E484:H |
| N...H | 3.759 | 0.0007 | G485:N | E484:HB2 |
| N...H | 4.175 | 0.0001 | G485:N | E484:H |
| N-C | 1.359 | 0.4099 | G485:N | E484:C |
| N-N | 3.687 | 0.0050 | V483:N | E484:N |
| O...H | 3.454 | 0.0003 | V483:HB | E484:O |
| O...H | 2.731 | 0.0016 | G485:HA3 | E484:O |
| O...H | 2.772 | 0.0024 | G485:HA2 | E484:O |
| O...H | 3.200 | 0.0084 | G485:H | E484:O |
| C-C | 4.373 | 0.0004 | E484:CD | Y489:C |
| C-C | 4.486 | 0.0003 | E484:CD | F490:CA |
| C-O | 3.637 | 0.0011 | E484:C | C488:O |
| H-C | 4.054 | 0.0001 | E484:HA | C488:CA |
| H-C | 4.487 | 0.0001 | E484:HB3 | C488:C |
| H-C | 4.474 | 0.0003 | E484:HG2 | Y489:CB |
| H-C | 3.988 | 0.0001 | E484:HG2 | F490:CA |
| H-H | 3.996 | 0.0002 | E484:HG3 | Y489:HA |
| H-H | 3.226 | 0.0002 | E484:HG3 | F490:H |
| O...H | 2.483 | 0.0012 | E484:HA | C488:O |
| C-O | 3.555 | 0.0025 | Y489:C | E484:OE1 |
| C-O | 3.543 | 0.0036 | F490:CA | E484:OE1 |
| H-C | 4.083 | 0.0001 | F490:HB3 | E484:CD |
| H-C | 4.292 | 0.0006 | F490:H | E484:CB |
| N-C | 3.571 | 0.0041 | F490:N | E484:CD |
| O...H | 2.581 | 0.0080 | K31:HZ2 | E484:OE1 |
| O...H | 2.844 | 0.0018 | K31:HE2 | E484:OE1 |
| O...H | 3.275 | 0.0004 | K31:HZ2 | E484:OE2 |
| O...H | 4.035 | 0.0001 | K31:HE3 | E484:OE1 |
| O...H | 2.762 | 0.0001 | Y489:HA | E484:OE1 |
| O...H | 1.557 | 0.0909 | F490:H | E484:OE1 |
| O...H | 3.709 | 0.0028 | F490:H | E484:OE2 |

**Table S10:** A484 with their bonding for OV interface model.

| A484 | | | | |
|---|---|---|---|---|
| C-C | 3.695 | 0.0138 | A484:C | G485:C |
| C-C | 3.817 | 0.0141 | A484:CA | G485:CA |
| C-O | 3.799 | 0.0010 | A484:C | V483:O |
| C-O | 4.046 | 0.0005 | A484:C | G485:O |
| C-O | 4.078 | 0.0006 | A484:CB | V483:O |
| H-C | 3.401 | 0.0001 | A484:H | V483:CG2 |
| H-C | 3.648 | 0.0009 | A484:HB3 | V483:C |
| H-C | 3.919 | 0.0006 | A484:H | V483:CG1 |
| H-C | 4.016 | 0.0009 | A484:HA | G485:CA |
| H-C | 4.072 | 0.0014 | A484:HA | V483:CA |
| O...H | 2.412 | 0.0061 | A484:HA | V483:O |
| O...H | 3.165 | 0.0080 | A484:H | V483:O |
| O...H | 4.027 | 0.0001 | A484:HB3 | V483:O |
| N-C | 1.362 | 0.4221 | A484:N | V483:C |
| N-C | 4.395 | 0.0003 | A484:N | V483:CG1 |
| N...H | 3.921 | 0.0011 | A484:N | G485:H |
| N...H | 4.091 | 0.0005 | A484:N | V483:H |
| N-N | 3.623 | 0.0050 | A484:N | G485:N |
| C-C | 3.408 | 0.0015 | V483:C | A484:C |
| C-C | 3.634 | 0.0095 | V483:C | A484:CB |
| C-C | 3.828 | 0.0126 | V483:CA | A484:CA |
| C-C | 4.357 | 0.0006 | V483:CB | A484:CA |
| C-O | 3.707 | 0.0001 | V483:C | A484:O |
| C-O | 4.269 | 0.0003 | G485:C | A484:O |
| H-C | 3.922 | 0.0009 | V483:HB | A484:CA |
| H-C | 4.103 | 0.0004 | V483:HA | A484:CA |
| H-C | 4.108 | 0.0002 | V483:HB | A484:C |
| H-C | 4.147 | 0.0007 | G485:HA3 | A484:CA |
| H-C | 4.254 | 0.0001 | G485:HA2 | A484:CA |
| H-H | 3.982 | 0.0001 | V483:HG22 | A484:H |
| H-H | 4.023 | 0.0001 | V483:HG13 | A484:H |
| H-H | 4.412 | 0.0001 | V483:HG12 | A484:H |
| O...H | 2.660 | 0.0026 | G485:HA3 | A484:O |
| O...H | 2.970 | 0.0014 | G485:HA2 | A484:O |
| O...H | 3.203 | 0.0083 | G485:H | A484:O |
| O...H | 3.498 | 0.0001 | V483:HB | A484:O |
| N-C | 1.353 | 0.4202 | G485:N | A484:C |
| N...H | 3.613 | 0.0005 | G485:N | A484:HB1 |
| N...H | 3.953 | 0.0008 | V483:N | A484:H |
| N...H | 3.981 | 0.0008 | G485:N | A484:H |
| N-N | 3.681 | 0.0053 | V483:N | A484:N |
| C-O | 3.644 | 0.0008 | A484:C | C488:O |
| O...H | 2.497 | 0.0014 | A484:HA | C488:O |
| O...H | 3.832 | 0.0001 | A484:HB3 | C488:O |
| H-H | 4.063 | 0.0001 | I472:HG23 | A484:HA |

**Table S11:** Q493 with their bonding for WT interface model.

| Q493 | | | | |
|---|---|---|---|---|
| C-C | 3.480 | 0.0034 | Q493:C | S494:C |
| C-C | 3.549 | 0.0072 | Q493:C | S494:CB |
| C-C | 3.835 | 0.0121 | Q493:CA | S494:CA |
| C-O | 3.733 | 0.0014 | Q493:CB | L492:O |
| C-O | 3.964 | 0.0006 | Q493:C | L492:O |
| C-O | 4.087 | 0.0002 | Q493:C | S494:O |
| C-O | 4.434 | 0.0001 | Q493:CG | L492:O |
| H-C | 3.327 | 0.0004 | Q493:H | L492:CB |
| H-C | 3.983 | 0.0020 | Q493:HA | L492:CA |
| H-C | 4.031 | 0.0007 | Q493:HA | S494:CA |
| H-C | 4.380 | 0.0001 | Q493:HG2 | L492:C |
| H-H | 4.070 | 0.0002 | Q493:HG3 | S494:H |
| O...H | 2.300 | 0.0074 | Q493:HA | L492:O |
| O...H | 3.189 | 0.0078 | Q493:H | L492:O |
| N-C | 1.350 | 0.4188 | Q493:N | L492:C |
| N...H | 3.653 | 0.0008 | Q493:N | L492:HB2 |
| N...H | 3.973 | 0.0009 | Q493:N | S494:H |
| N...H | 3.976 | 0.0006 | Q493:N | L492:H |
| N-N | 3.635 | 0.0052 | Q493:N | S494:N |
| C-C | 3.419 | 0.0021 | L492:C | Q493:CB |
| C-C | 3.505 | 0.0082 | L492:C | Q493:C |
| C-C | 3.819 | 0.0134 | L492:CA | Q493:CA |
| C-O | 3.768 | 0.0002 | L492:C | Q493:O |
| C-O | 3.788 | 0.0008 | S494:C | Q493:O |
| C-O | 3.996 | 0.0012 | S494:CB | Q493:O |
| H-C | 3.495 | 0.0007 | S494:HB2 | Q493:C |
| H-C | 3.928 | 0.0001 | S494:HB3 | Q493:C |
| H-C | 3.985 | 0.0012 | L492:HA | Q493:CA |
| H-C | 4.014 | 0.0016 | S494:HA | Q493:CA |
| O...H | 2.324 | 0.0062 | S494:HA | Q493:O |
| O...H | 3.180 | 0.0076 | S494:H | Q493:O |
| O...H | 3.895 | 0.0001 | S494:HB2 | Q493:O |
| N-C | 1.355 | 0.4486 | S494:N | Q493:C |
| N...H | 3.352 | 0.0002 | S494:N | Q493:HB3 |
| N...H | 3.825 | 0.0012 | L492:N | Q493:H |
| N...H | 3.969 | 0.0009 | S494:N | Q493:H |
| N-N | 3.540 | 0.0046 | L492:N | Q493:N |
| C-O | 3.577 | 0.0017 | Q493:CD | Y453:OH |
| C-O | 3.676 | 0.0008 | Q493:CD | E35:OE1 |
| C-O | 3.895 | 0.0003 | Q493:CD | H34:O |
| C-O | 3.941 | 0.0007 | Q493:CG | Y453:OH |
| C-O | 4.145 | 0.0011 | Q493:CA | Y453:O |
| C-O | 4.240 | 0.0003 | Q493:C | Y453:O |
| H-C | 3.838 | 0.0006 | Q493:HE21 | H34:CA |
| H-C | 3.959 | 0.0001 | Q493:H | Y453:CA |
| H-C | 4.000 | 0.0002 | Q493:HB2 | Y453:CZ |
| H-C | 4.309 | 0.0002 | Q493:HB2 | Y453:CE1 |
| H-C | 4.374 | 0.0001 | Q493:HE21 | E35:CD |
| H-C | 4.417 | 0.0001 | Q493:HB2 | L455:CG |
| H-C | 4.434 | 0.0001 | Q493:H | P491:C |
| O...H | 1.867 | 0.0533 | Q493:HE22 | E35:OE1 |
| O...H | 2.045 | 0.0212 | Q493:H | Y453:O |
| O...H | 2.165 | 0.0149 | Q493:HE21 | H34:O |
| O...H | 3.482 | 0.0002 | Q493:HE22 | H34:O |
| O...H | 3.534 | 0.0005 | Q493:HE21 | E35:OE1 |
| O...H | 3.900 | 0.0001 | Q493:HB2 | Y453:OH |
| O...H | 3.964 | 0.0001 | Q493:HG3 | Y453:OH |
| O...H | 4.100 | 0.0010 | Q493:HE22 | E35:OE2 |
| N-C | 3.921 | 0.0018 | Q493:NE2 | E35:CD |
| N-C | 4.152 | 0.0011 | Q493:N | Y453:C |
| N-C | 4.259 | 0.0004 | Q493:N | P491:C |
| N-C | 4.305 | 0.0003 | Q493:NE2 | E35:CG |
| N...H | 3.790 | 0.0022 | Q493:NE2 | Y453:HH |
| N...H | 4.326 | 0.0001 | Q493:N | R454:HA |
| C-C | 4.205 | 0.0002 | H34:C | Q493:CD |

**Table S12:** R493 with their bonding for OV interface model.

| R493 | | | | |
|---|---|---|---|---|
| C-C | 3.665 | 0.0121 | R493:C | S494:CB |
| C-C | 3.824 | 0.0127 | R493:CA | S494:CA |
| C-C | 4.260 | 0.0001 | R493:CD | S494:CA |
| C-O | 3.520 | 0.0011 | R493:CD | S494:O |
| C-O | 3.660 | 0.0021 | R493:CZ | S494:O |
| C-O | 3.863 | 0.0003 | R493:C | S494:O |
| C-O | 4.135 | 0.0004 | R493:C | L492:O |
| C-O | 4.413 | 0.0002 | R493:CG | L492:O |
| H-C | 3.490 | 0.0001 | R493:HB3 | L492:C |
| H-C | 3.971 | 0.0010 | R493:HA | S494:CA |
| H-C | 4.034 | 0.0013 | R493:HA | L492:CA |
| H-C | 4.053 | 0.0008 | R493:HG2 | S494:CA |
| H-C | 4.070 | 0.0002 | R493:HB2 | L492:CA |
| H-C | 4.438 | 0.0002 | R493:HG2 | S494:CB |
| H-H | 3.778 | 0.0004 | R493:HG3 | S494:H |
| H-H | 4.305 | 0.0001 | R493:HE | S494:HA |
| N...H | 3.583 | 0.0008 | R493:N | L492:HB2 |
| N...H | 3.907 | 0.0009 | R493:N | S494:H |
| N...H | 4.006 | 0.0005 | R493:N | L492:H |
| N-C | 1.349 | 0.4340 | R493:N | L492:C |
| N-C | 3.667 | 0.0024 | R493:NE | S494:C |
| N-C | 4.046 | 0.0006 | R493:NE | S494:CA |
| N-N | 3.626 | 0.0046 | R493:N | S494:N |
| N-O | 3.739 | 0.0001 | R493:NH1 | S494:O |
| O...H | 1.699 | 0.0605 | R493:HE | S494:O |
| O...H | 2.391 | 0.0045 | R493:HA | L492:O |
| O...H | 2.652 | 0.0002 | R493:HB2 | L492:O |
| O...H | 3.112 | 0.0023 | R493:HD3 | S494:O |
| O...H | 3.199 | 0.0080 | R493:H | L492:O |
| O...H | 3.905 | 0.0001 | R493:HB3 | L492:O |
| C-C | 3.640 | 0.0124 | L492:C | R493:C |
| C-C | 3.804 | 0.0140 | L492:CA | R493:CA |
| C-O | 3.401 | 0.0004 | S494:C | R493:O |
| C-O | 4.020 | 0.0005 | L492:C | R493:O |
| C-O | 4.215 | 0.0009 | S494:CB | R493:O |
| H-C | 3.186 | 0.0011 | S494:H | R493:CB |
| H-C | 3.707 | 0.0009 | S494:HB2 | R493:C |
| H-C | 3.987 | 0.0011 | L492:HA | R493:CA |
| H-C | 3.988 | 0.0004 | S494:HB3 | R493:C |
| H-C | 4.036 | 0.0010 | S494:HA | R493:CA |
| H-C | 4.354 | 0.0001 | S494:H | R493:CZ |
| N...H | 3.781 | 0.0003 | S494:N | R493:HB3 |
| N...H | 3.885 | 0.0011 | L492:N | R493:H |
| N...H | 3.924 | 0.0012 | S494:N | R493:H |
| N...H | 4.284 | 0.0003 | S494:N | R493:HG3 |
| N...H | 4.366 | 0.0003 | S494:N | R493:HD2 |
| N-C | 1.357 | 0.4531 | S494:N | R493:C |
| N-N | 3.583 | 0.0049 | L492:N | R493:N |
| O...H | 2.526 | 0.0030 | S494:HA | R493:O |
| O...H | 3.163 | 0.0086 | S494:H | R493:O |
| C-O | 3.861 | 0.0018 | R493:CZ | E35:OE1 |
| C-O | 4.078 | 0.0013 | R493:CA | Y453:O |
| C-O | 4.155 | 0.0003 | R493:C | Y453:O |
| C-O | 4.207 | 0.0002 | R493:CD | E35:OE1 |
| H-C | 3.287 | 0.0027 | R493:HH22 | H34:CA |
| H-C | 3.635 | 0.0012 | R493:HH21 | H34:C |
| H-C | 3.759 | 0.0004 | R493:HH22 | E35:CB |
| H-C | 4.184 | 0.0002 | R493:HH12 | D38:CA |
| H-C | 4.197 | 0.0001 | R493:HH12 | H34:CA |
| H-C | 4.246 | 0.0001 | R493:HE | Y453:CE1 |
| H-C | 4.412 | 0.0002 | R493:HH12 | D38:CG |
| N...H | 4.084 | 0.0001 | R493:NH1 | D38:HB3 |
| N...H | 4.271 | 0.0001 | R493:NH2 | D38:HB2 |
| N...H | 4.424 | 0.0001 | R493:N | R454:HA |
| N-C | 3.707 | 0.0016 | R493:NH2 | E35:CD |

**Table S11:** Q493 with their bonding for WT interface model.

| | | | | |
|---|---|---|---|---|
| Q493 | | | | |
| C-C | 4.306 | 0.0007 | Y453:CZ | Q493:CD |
| C-C | 4.309 | 0.0004 | Y453:CE1 | Q493:CD |
| C-C | 4.483 | 0.0005 | Y453:CZ | Q493:CG |
| C-C | 4.485 | 0.0008 | L452:CA | Q493:C |
| C-O | 3.372 | 0.0017 | Y453:CZ | Q493:OE1 |
| C-O | 3.697 | 0.0010 | L452:C | Q493:O |
| C-O | 4.085 | 0.0012 | Y453:CA | Q493:O |
| C-O | 4.173 | 0.0002 | Y453:C | Q493:O |
| C-O | 4.419 | 0.0002 | L452:CG | Q493:O |
| C-O | 4.450 | 0.0001 | H34:C | Q493:OE1 |
| H-C | 3.686 | 0.0002 | Y453:HE1 | Q493:CD |
| H-C | 4.002 | 0.0001 | Y453:H | Q493:CA |
| H-C | 4.026 | 0.0001 | H34:HB3 | Q493:CG |
| H-C | 4.054 | 0.0002 | L452:HD21 | Q493:C |
| H-C | 4.160 | 0.0001 | Y453:HE1 | Q493:CA |
| H-C | 4.231 | 0.0001 | Y495:H | Q493:C |
| H-C | 4.243 | 0.0002 | Y453:HE1 | Q493:CG |
| H-C | 4.413 | 0.0002 | Y453:HH | Q493:CA |
| H-H | 3.919 | 0.0001 | H34:HB3 | Q493:HE22 |
| H-H | 3.977 | 0.0001 | Y453:HH | Q493:HE21 |
| H-H | 4.222 | 0.0001 | H34:HA | Q493:HE21 |
| H-H | 4.248 | 0.0001 | L455:HB2 | Q493:HG3 |
| H-H | 4.294 | 0.0001 | L455:HD22 | Q493:HG3 |
| H-H | 4.297 | 0.0001 | Y453:HH | Q493:HG2 |
| H-H | 4.462 | 0.0001 | Y453:HE1 | Q493:HB2 |
| O...H | 1.643 | 0.0887 | Y453:HH | Q493:OE1 |
| O...H | 2.022 | 0.0220 | Y453:H | Q493:O |
| O...H | 2.453 | 0.0029 | L452:HA | Q493:O |
| O...H | 3.256 | 0.0001 | L452:HB3 | Q493:O |
| N-C | 4.205 | 0.0003 | Y495:N | Q493:C |
| N-C | 4.211 | 0.0016 | Y453:N | Q493:C |
| N...H | 4.138 | 0.0003 | R454:N | Q493:H |
| N-N | 3.822 | 0.0003 | E35:N | Q493:NE2 |

**Table S12:** R493 with their bonding for OV interface model.

| | | | | |
|---|---|---|---|---|
| R493 | | | | |
| N-C | 4.004 | 0.0001 | R493:NH2 | E35:CG |
| N-C | 4.021 | 0.0017 | R493:NH1 | H34:C |
| N-C | 4.140 | 0.0017 | R493:N | Y453:C |
| N-C | 4.288 | 0.0001 | R493:NH2 | E35:CB |
| N-C | 4.327 | 0.0003 | R493:N | P491:C |
| N-C | 4.498 | 0.0001 | R493:NE | Y453:CE1 |
| N-O | 4.273 | 0.0004 | R493:NE | E35:OE1 |
| N-O | 4.423 | 0.0001 | R493:NE | Y453:OH |
| O...H | 1.804 | 0.0616 | R493:HH21 | E35:OE1 |
| O...H | 1.809 | 0.0407 | R493:HH22 | H34:O |
| O...H | 1.979 | 0.0253 | R493:H | Y453:O |
| O...H | 2.136 | 0.0135 | R493:HH12 | H34:O |
| O...H | 3.176 | 0.0001 | R493:HH22 | E35:OE1 |
| O...H | 3.548 | 0.0010 | R493:HH21 | H34:O |
| O...H | 3.713 | 0.0004 | R493:HH11 | H34:O |
| O...H | 4.034 | 0.0012 | R493:HH21 | E35:OE2 |
| O...H | 4.277 | 0.0001 | R493:HD2 | E35:OE1 |
| O...H | 4.349 | 0.0001 | R493:HH22 | E35:O |
| C-C | 3.972 | 0.0018 | H34:C | R493:CZ |
| C-O | 3.669 | 0.0012 | L452:C | R493:O |
| C-O | 3.909 | 0.0018 | Y453:CA | R493:O |
| C-O | 4.025 | 0.0003 | Y453:C | R493:O |
| H-C | 3.841 | 0.0007 | E35:HA | R493:CZ |
| H-C | 3.890 | 0.0002 | L452:HD21 | R493:C |
| H-C | 3.908 | 0.0007 | D38:HB2 | R493:CZ |
| H-C | 3.964 | 0.0002 | Y453:H | R493:CA |
| H-C | 4.214 | 0.0001 | Y495:H | R493:C |
| H-H | 3.092 | 0.0003 | E35:HA | R493:HH21 |
| H-H | 3.297 | 0.0001 | D38:HB2 | R493:HH11 |
| H-H | 3.483 | 0.0001 | H34:HB3 | R493:HH21 |
| H-H | 3.592 | 0.0001 | D38:HB3 | R493:HH12 |
| H-H | 3.685 | 0.0001 | Y453:HE1 | R493:HD2 |
| H-H | 3.788 | 0.0003 | H34:HA | R493:HH22 |
| N...H | 3.892 | 0.0017 | Y495:N | R493:HE |
| N...H | 4.137 | 0.0002 | E35:N | R493:HH12 |
| N...H | 4.176 | 0.0004 | R454:N | R493:H |
| N-C | 4.121 | 0.0026 | Y453:N | R493:C |
| N-C | 4.144 | 0.0003 | Y495:N | R493:C |
| N-N | 3.421 | 0.0019 | E35:N | R493:NH2 |
| N-N | 4.432 | 0.0003 | Y453:N | R493:N |
| O...H | 1.895 | 0.0288 | Y453:H | R493:O |
| O...H | 2.548 | 0.0018 | L452:HA | R493:O |
| O...H | 3.521 | 0.0001 | L452:HB3 | R493:O |
| O...H | 3.754 | 0.0001 | Y495:H | R493:O |

**Table S13:** G496 with their bonding for WT interface model.

| G496 | | | | |
|---|---|---|---|---|
| C-C | 3.687 | 0.0117 | G496:C | F497:CB |
| C-C | 3.813 | 0.0144 | G496:CA | F497:CA |
| C-O | 4.030 | 0.0001 | G496:C | F497:O |
| H-C | 3.339 | 0.0116 | G496:HA3 | Y495:C |
| H-C | 3.851 | 0.0012 | G496:H | Y495:CB |
| H-C | 3.975 | 0.0010 | G496:HA2 | F497:CA |
| H-C | 4.167 | 0.0005 | G496:HA2 | Y495:CA |
| H-H | 3.482 | 0.0011 | G496:HA3 | F497:H |
| O...H | 2.604 | 0.0043 | G496:HA2 | Y495:O |
| O...H | 3.142 | 0.0078 | G496:H | Y495:O |
| O...H | 3.829 | 0.0003 | G496:HA3 | Y495:O |
| N-C | 1.360 | 0.4246 | G496:N | Y495:C |
| N-C | 3.543 | 0.0037 | G496:N | Y495:CB |
| N...H | 3.391 | 0.0006 | G496:N | Y495:HB3 |
| N...H | 4.035 | 0.0004 | G496:N | Y495:HB2 |
| C-C | 3.822 | 0.0142 | Y495:CA | G496:CA |
| C-O | 3.618 | 0.0008 | F497:C | G496:O |
| C-O | 4.051 | 0.0002 | Y495:C | G496:O |
| C-O | 4.211 | 0.0006 | F497:CB | G496:O |
| H-C | 3.918 | 0.0012 | Y495:HA | G496:CA |
| H-C | 4.004 | 0.0002 | F497:HD1 | G496:C |
| H-C | 4.021 | 0.0004 | F497:HB2 | G496:C |
| H-C | 4.068 | 0.0009 | F497:HA | G496:CA |
| H-H | 3.847 | 0.0001 | Y495:HB3 | G496:H |
| O...H | 2.472 | 0.0041 | F497:HA | G496:O |
| O...H | 3.157 | 0.0077 | F497:H | G496:O |
| N-C | 1.350 | 0.4694 | F497:N | G496:C |
| N...H | 3.288 | 0.0047 | F497:N | G496:HA3 |
| N...H | 3.307 | 0.0008 | Y495:N | G496:H |
| N-N | 3.370 | 0.0033 | Y495:N | G496:N |
| C-O | 4.142 | 0.0001 | G496:C | D38:OD1 |
| C-O | 4.326 | 0.0008 | G496:CA | S494:O |
| H-C | 4.254 | 0.0004 | G496:HA3 | D38:CB |
| H-C | 4.360 | 0.0006 | G496:HA3 | K353:CE |
| O...H | 2.322 | 0.0066 | G496:HA3 | D38:OD1 |
| O...H | 2.494 | 0.0064 | G496:H | S494:O |
| O...H | 3.901 | 0.0001 | G496:HA2 | D38:OD1 |
| O...H | 3.979 | 0.0003 | G496:H | D38:OD1 |
| O...H | 4.497 | 0.0001 | G496:HA3 | S494:O |
| N-C | 4.434 | 0.0001 | G496:N | Y449:CE1 |
| N...H | 3.573 | 0.0006 | G496:N | K353:HZ1 |
| N...H | 3.956 | 0.0002 | G496:N | K353:HZ3 |
| N-O | 4.187 | 0.0002 | G496:N | D38:OD1 |
| C-O | 3.886 | 0.0019 | K353:CE | G496:O |
| C-O | 4.259 | 0.0005 | N501:CG | G496:O |
| H-C | 3.724 | 0.0005 | K353:HZ3 | G496:C |
| H-C | 3.815 | 0.0002 | Q498:H | G496:C |
| H-C | 4.395 | 0.0001 | N448:H | G496:C |
| H-H | 3.710 | 0.0003 | K353:HZ3 | G496:H |
| H-H | 4.036 | 0.0003 | K353:HZ1 | G496:HA2 |
| H-H | 4.233 | 0.0001 | K353:HZ3 | G496:HA2 |
| O...H | 1.793 | 0.1000 | K353:HZ1 | G496:O |
| O...H | 2.280 | 0.0126 | N501:HD21 | G496:O |
| O...H | 3.725 | 0.0003 | N501:HD22 | G496:O |
| N-C | 3.535 | 0.0009 | K353:NZ | G496:C |
| N-C | 3.905 | 0.0004 | Q498:N | G496:C |
| N-C | 4.264 | 0.0006 | N501:ND2 | G496:C |

**Table S14:** S496 with their bonding for OV interface model.

| S496 | | | | |
|---|---|---|---|---|
| C-C | 3.471 | 0.0001 | S496:C | F497:C |
| C-C | 3.656 | 0.0107 | S496:C | F497:CB |
| C-C | 3.820 | 0.0136 | S496:CA | F497:CA |
| C-O | 3.351 | 0.0001 | S496:C | Y495:O |
| C-O | 4.000 | 0.0002 | S496:C | F497:O |
| C-O | 4.236 | 0.0007 | S496:CB | Y495:O |
| H-C | 3.764 | 0.0008 | S496:HB2 | Y495:C |
| H-C | 3.929 | 0.0012 | S496:HA | F497:CA |
| H-C | 3.931 | 0.0014 | S496:H | Y495:CB |
| H-C | 3.984 | 0.0005 | S496:HB3 | Y495:C |
| H-C | 4.065 | 0.0010 | S496:HA | Y495:CA |
| N...H | 3.122 | 0.0002 | S496:N | F497:H |
| N...H | 3.671 | 0.0015 | S496:N | Y495:HB3 |
| N...H | 3.786 | 0.0001 | S496:N | Y495:H |
| N...H | 4.298 | 0.0001 | S496:N | Y495:HD1 |
| N-C | 1.362 | 0.4151 | S496:N | Y495:C |
| N-C | 3.699 | 0.0057 | S496:N | Y495:CB |
| N-N | 3.234 | 0.0003 | S496:N | F497:N |
| O...H | 2.561 | 0.0045 | S496:HA | Y495:O |
| O...H | 3.165 | 0.0085 | S496:H | Y495:O |
| C-C | 3.675 | 0.0129 | Y495:C | S496:CB |
| C-C | 3.813 | 0.0138 | Y495:CA | S496:CA |
| C-O | 3.761 | 0.0006 | F497:C | S496:O |
| C-O | 4.084 | 0.0001 | Y495:C | S496:O |
| C-O | 4.205 | 0.0006 | F497:CB | S496:O |
| H-C | 3.852 | 0.0017 | F497:H | S496:CB |
| H-C | 3.958 | 0.0012 | Y495:HA | S496:CA |
| H-C | 4.044 | 0.0002 | F497:HB2 | S496:C |
| H-C | 4.049 | 0.0011 | F497:HA | S496:CA |
| H-C | 4.146 | 0.0001 | F497:HD1 | S496:C |
| H-H | 4.098 | 0.0001 | Y495:HD1 | S496:H |
| N...H | 4.140 | 0.0004 | F497:N | S496:HB3 |
| N-C | 1.358 | 0.4463 | F497:N | S496:C |
| N-C | 3.656 | 0.0052 | F497:N | S496:CB |
| O...H | 2.477 | 0.0053 | F497:HA | S496:O |
| O...H | 3.160 | 0.0080 | F497:H | S496:O |
| C-C | 4.255 | 0.0014 | S496:C | Y501:CE2 |
| C-O | 4.123 | 0.0013 | S496:CA | S494:O |
| C-O | 4.174 | 0.0001 | S496:CB | D38:OD2 |
| C-O | 4.498 | 0.0001 | S496:CB | Y449:OH |
| H-C | 3.855 | 0.0003 | S496:HB2 | Y449:CZ |
| H-C | 3.861 | 0.0033 | S496:HG | D38:CB |
| H-C | 4.073 | 0.0009 | S496:H | S494:CA |
| H-C | 4.090 | 0.0002 | S496:HA | Y449:CD1 |
| H-C | 4.265 | 0.0001 | S496:HA | Y449:CG |
| H-C | 4.283 | 0.0001 | S496:HG | Y449:CD1 |
| H-C | 4.329 | 0.0002 | S496:HB3 | Y449:CE1 |
| H-C | 4.436 | 0.0001 | S496:HA | Y449:CE1 |
| H-C | 4.474 | 0.0002 | S496:HB3 | K353:CE |
| H-H | 3.247 | 0.0005 | S496:HG | R498:HH11 |
| H-H | 3.881 | 0.0002 | S496:HB3 | R498:HH12 |
| N...H | 4.218 | 0.0001 | S496:N | Y449:HA |
| N...H | 4.287 | 0.0001 | S496:N | K353:HZ2 |
| O...H | 1.751 | 0.0658 | S496:HG | D38:OD1 |
| O...H | 2.203 | 0.0090 | S496:H | S494:O |
| O...H | 2.833 | 0.0016 | S496:HB3 | D38:OD1 |
| C-O | 3.935 | 0.0003 | Y501:CZ | S496:O |
| C-O | 4.217 | 0.0012 | K353:CE | S496:O |
| C-O | 4.305 | 0.0006 | Y501:CD2 | S496:O |
| C-O | 4.360 | 0.0001 | Y449:CG | S496:OG |
| C-O | 4.406 | 0.0001 | Y449:CD2 | S496:OG |
| H-C | 3.919 | 0.0011 | R498:HH12 | S496:CB |
| H-C | 4.150 | 0.0002 | Y449:HH | S496:CB |
| H-C | 4.166 | 0.0001 | K353:HZ1 | S496:C |
| H-C | 4.223 | 0.0001 | Y449:HA | S496:CB |
| H-C | 4.264 | 0.0001 | K353:HZ3 | S496:C |
| H-C | 4.359 | 0.0001 | R498:H | S496:C |
| H-C | 4.414 | 0.0001 | N448:H | S496:C |
| H-H | 3.436 | 0.0001 | K353:HZ2 | S496:HG |
| H-H | 3.772 | 0.0001 | Y449:HA | S496:HB2 |
| H-H | 3.918 | 0.0003 | K353:HZ1 | S496:HB3 |
| H-H | 3.981 | 0.0001 | Y449:HE1 | S496:HB3 |
| N...H | 4.496 | 0.0001 | R498:NH1 | S496:HB3 |
| N-C | 4.095 | 0.0002 | K353:NZ | S496:CB |
| N-C | 4.097 | 0.0013 | K353:NZ | S496:C |
| N-C | 4.306 | 0.0002 | R498:N | S496:C |
| N-C | 4.432 | 0.0006 | R498:NH1 | S496:CB |
| O...H | 2.108 | 0.0392 | K353:HZ2 | S496:O |
| O...H | 2.213 | 0.0016 | Y501:HE2 | S496:O |
| O...H | 4.358 | 0.0002 | Y501:HD2 | S496:O |
| O...H | 4.426 | 0.0001 | K353:HE2 | S496:O |

**Table S15:** Q498 with their bonding for WT interface model.

| Q498 | | | | |
|---|---|---|---|---|
| C-C | 3.662 | 0.0134 | Q498:C | P499:CB |
| C-C | 3.676 | 0.0120 | Q498:C | P499:CG |
| C-C | 3.870 | 0.0120 | Q498:CA | P499:CA |
| C-C | 4.446 | 0.0006 | Q498:CA | P499:CG |
| C-C | 4.457 | 0.0001 | Q498:CG | P499:CD |
| C-O | 4.022 | 0.0007 | Q498:CB | F497:O |
| C-O | 4.027 | 0.0005 | Q498:C | F497:O |
| C-O | 4.235 | 0.0001 | Q498:C | P499:O |
| H-C | 3.608 | 0.0018 | Q498:H | F497:CB |
| H-C | 3.795 | 0.0002 | Q498:HB2 | F497:C |
| H-C | 4.056 | 0.0002 | Q498:HG3 | F497:C |
| H-C | 4.094 | 0.0012 | Q498:HA | F497:CA |
| H-C | 4.100 | 0.0005 | Q498:HA | P499:CA |
| H-C | 4.180 | 0.0008 | Q498:HA | P499:CG |
| H-C | 4.268 | 0.0002 | Q498:HB3 | P499:CG |
| H-C | 4.424 | 0.0002 | Q498:HB3 | P499:C |
| H-H | 3.899 | 0.0001 | Q498:HB3 | P499:HD3 |
| H-H | 3.941 | 0.0001 | Q498:HG2 | P499:HD2 |
| H-H | 4.024 | 0.0001 | Q498:HB2 | P499:HD2 |
| H-H | 4.095 | 0.0001 | Q498:HB3 | P499:HG2 |
| H-H | 4.181 | 0.0001 | Q498:HG3 | P499:HD3 |
| H-H | 4.385 | 0.0001 | Q498:HA | P499:HA |
| N...H | 3.987 | 0.0004 | Q498:N | F497:HB2 |
| N...H | 4.233 | 0.0002 | Q498:N | P499:HD3 |
| N...H | 4.240 | 0.0002 | Q498:N | P499:HD2 |
| N...H | 4.374 | 0.0001 | Q498:N | F497:HD1 |
| N-C | 1.338 | 0.4593 | Q498:N | F497:C |
| N-C | 3.475 | 0.0024 | Q498:N | F497:CB |
| N-C | 4.271 | 0.0011 | Q498:N | P499:CD |
| N-N | 3.584 | 0.0045 | Q498:N | P499:N |
| O...H | 2.497 | 0.0046 | Q498:HA | F497:O |
| O...H | 3.190 | 0.0083 | Q498:H | F497:O |
| C-C | 3.496 | 0.0066 | F497:C | Q498:C |
| C-C | 3.576 | 0.0054 | F497:C | Q498:CB |
| C-C | 3.825 | 0.0144 | F497:CA | Q498:CA |
| C-C | 4.028 | 0.0006 | F497:C | Q498:CG |
| C-O | 3.658 | 0.0097 | P499:CD | Q498:O |
| C-O | 3.820 | 0.0006 | F497:C | Q498:O |
| C-O | 4.227 | 0.0011 | P499:CB | Q498:O |
| H-C | 3.877 | 0.0015 | F497:HA | Q498:CA |
| H-C | 4.023 | 0.0003 | P499:HG2 | Q498:C |
| H-C | 4.055 | 0.0002 | P499:HB3 | Q498:C |
| H-C | 4.124 | 0.0006 | P499:HA | Q498:CA |
| H-H | 3.450 | 0.0002 | F497:HB3 | Q498:H |
| H-H | 3.792 | 0.0001 | F497:HD1 | Q498:H |
| N...H | 3.511 | 0.0017 | F497:N | Q498:H |
| N...H | 3.888 | 0.0002 | P499:N | Q498:HB2 |
| N...H | 3.911 | 0.0011 | P499:N | Q498:H |
| N-C | 1.358 | 0.4974 | P499:N | Q498:C |
| N-N | 3.393 | 0.0026 | F497:N | Q498:N |
| O...H | 2.669 | 0.0009 | P499:HA | Q498:O |
| O...H | 3.981 | 0.0001 | F497:H | Q498:OE1 |
| O...H | 4.009 | 0.0005 | P499:HD2 | Q498:O |
| O...H | 4.082 | 0.0003 | P499:HD3 | Q498:O |
| C-C | 4.318 | 0.0004 | Q498:CA | N501:CG |
| C-C | 4.403 | 0.0004 | Q498:C | T500:CA |
| C-O | 3.989 | 0.0002 | Q498:CA | N501:OD1 |
| C-O | 4.184 | 0.0002 | Q498:C | S443:OG |
| C-O | 4.342 | 0.0006 | Q498:CD | Q42:OE1 |
| C-O | 4.419 | 0.0001 | Q498:CB | K444:O |
| H-C | 3.782 | 0.0006 | Q498:HE22 | Q42:CG |
| H-C | 3.815 | 0.0002 | Q498:H | G496:C |
| H-C | 4.017 | 0.0001 | Q498:HG2 | Y41:CD2 |
| H-C | 4.061 | 0.0001 | Q498:HG3 | G446:CA |
| H-C | 4.151 | 0.0004 | Q498:HB3 | T500:CB |

**Table S16:** R498 with their bonding for OV interface model.

| R498 | | | | |
|---|---|---|---|---|
| C-C | 3.664 | 0.0131 | R498:C | P499:CB |
| C-C | 3.684 | 0.0121 | R498:C | P499:CG |
| C-C | 3.877 | 0.0121 | R498:CA | P499:CA |
| C-C | 4.393 | 0.0003 | R498:CG | P499:CD |
| C-C | 4.423 | 0.0006 | R498:CA | P499:CG |
| C-O | 3.903 | 0.0005 | R498:CB | F497:O |
| C-O | 4.017 | 0.0002 | R498:CD | F497:O |
| C-O | 4.099 | 0.0005 | R498:C | F497:O |
| H-C | 3.383 | 0.0009 | R498:H | F497:CB |
| H-C | 3.888 | 0.0001 | R498:HB2 | F497:C |
| H-C | 3.931 | 0.0004 | R498:HB3 | P499:CG |
| H-C | 4.068 | 0.0012 | R498:HA | F497:CA |
| H-C | 4.140 | 0.0004 | R498:HA | P499:CA |
| H-C | 4.204 | 0.0004 | R498:HB2 | P499:CD |
| H-C | 4.224 | 0.0007 | R498:HA | P499:CG |
| H-C | 4.437 | 0.0001 | R498:HB3 | P499:C |
| H-H | 3.552 | 0.0005 | R498:HB3 | P499:HD3 |
| H-H | 3.668 | 0.0002 | R498:HG2 | P499:HD2 |
| H-H | 3.690 | 0.0004 | R498:HB2 | P499:HD2 |
| H-H | 3.785 | 0.0002 | R498:HB3 | P499:HG2 |
| H-H | 4.407 | 0.0001 | R498:HA | P499:HA |
| N...H | 3.738 | 0.0009 | R498:N | F497:HB2 |
| N...H | 4.257 | 0.0002 | R498:N | P499:HD2 |
| N...H | 4.320 | 0.0002 | R498:N | P499:HD3 |
| N-C | 1.347 | 0.4587 | R498:N | F497:C |
| N-C | 4.325 | 0.0011 | R498:N | P499:CD |
| N-N | 3.646 | 0.0042 | R498:N | P499:N |
| O...H | 2.485 | 0.0050 | R498:HA | F497:O |
| O...H | 3.192 | 0.0082 | R498:H | F497:O |
| C-C | 3.530 | 0.0068 | F497:C | R498:C |
| C-C | 3.573 | 0.0057 | F497:C | R498:CB |
| C-C | 3.839 | 0.0002 | F497:C | R498:CG |
| C-C | 3.842 | 0.0130 | F497:CA | R498:CA |
| C-O | 3.665 | 0.0094 | P499:CD | R498:O |
| C-O | 3.768 | 0.0007 | F497:C | R498:O |
| C-O | 4.253 | 0.0009 | P499:CB | R498:O |
| H-C | 3.974 | 0.0011 | F497:HA | R498:CA |
| H-C | 4.040 | 0.0003 | P499:HB3 | R498:C |
| H-C | 4.059 | 0.0002 | P499:HG2 | R498:C |
| H-C | 4.140 | 0.0006 | P499:HA | R498:CA |
| H-H | 3.937 | 0.0001 | F497:HD1 | R498:H |
| N...H | 3.632 | 0.0002 | P499:N | R498:HB2 |
| N...H | 3.801 | 0.0012 | F497:N | R498:H |
| N...H | 3.823 | 0.0019 | P499:N | R498:H |
| N-C | 1.358 | 0.4864 | P499:N | R498:C |
| N-N | 3.588 | 0.0041 | F497:N | R498:N |
| O...H | 2.665 | 0.0008 | P499:HA | R498:O |
| O...H | 4.043 | 0.0005 | P499:HD3 | R498:O |
| O...H | 4.045 | 0.0003 | P499:HD2 | R498:O |
| C-C | 4.455 | 0.0001 | R498:CG | Y501:CZ |
| C-C | 4.464 | 0.0002 | R498:CA | Y501:CZ |
| C-C | 4.481 | 0.0005 | R498:CA | Y501:CE2 |
| C-O | 3.628 | 0.0007 | R498:CZ | S446:O |
| C-O | 3.858 | 0.0014 | R498:CZ | Y449:OH |
| C-O | 3.902 | 0.0014 | R498:CZ | D38:OD1 |
| C-O | 3.905 | 0.0014 | R498:CZ | Y501:OH |
| C-O | 4.101 | 0.0002 | R498:C | S443:OG |
| C-O | 4.262 | 0.0001 | R498:CB | K444:O |
| C-O | 4.359 | 0.0007 | R498:CG | Y501:OH |
| C-O | 4.426 | 0.0002 | R498:CB | Y501:OH |
| C-O | 4.426 | 0.0002 | R498:C | N439:OD1 |
| H-C | 3.733 | 0.0003 | R498:HH21 | Y449:CZ |
| H-C | 3.855 | 0.0014 | R498:HH22 | Y449:CE1 |
| H-C | 3.919 | 0.0011 | R498:HH12 | S496:CB |
| H-C | 3.955 | 0.0002 | R498:HH22 | D38:CG |
| H-C | 3.983 | 0.0001 | R498:HH21 | Q42:CG |
| H-C | 3.986 | 0.0016 | R498:HH21 | S446:CA |
| H-C | 4.116 | 0.0005 | R498:HH11 | Y501:CE1 |
| H-C | 4.123 | 0.0013 | R498:HH12 | D38:CB |
| H-C | 4.175 | 0.0001 | R498:HD3 | Y41:CZ |
| H-C | 4.213 | 0.0001 | R498:HD3 | Y41:CG |
| H-C | 4.235 | 0.0001 | R498:HH11 | Y41:CG |
| H-C | 4.259 | 0.0001 | R498:HB3 | T500:CB |
| H-C | 4.265 | 0.0001 | R498:HH12 | Y449:CZ |
| H-C | 4.272 | 0.0003 | R498:HH22 | Y449:CD2 |
| H-C | 4.289 | 0.0001 | R498:HA | K444:CA |

**Table S15:** Q498 with their bonding for WT interface model.

| Q498 | | | | |
|---|---|---|---|---|
| H-C | 4.219 | 0.0001 | Q498:HG3 | V445:C |
| H-C | 4.377 | 0.0001 | Q498:HB3 | T500:CA |
| H-C | 4.405 | 0.0001 | Q498:HA | K444:CA |
| H-C | 4.458 | 0.0002 | Q498:HB2 | N501:CB |
| H-H | 3.210 | 0.0005 | Q498:H | N501:HD22 |
| H-H | 3.678 | 0.0001 | Q498:HB2 | T500:H |
| N...H | 3.704 | 0.0001 | Q498:NE2 | Y41:HE2 |
| N...H | 4.107 | 0.0008 | Q498:N | N448:H |
| N-C | 3.905 | 0.0004 | Q498:N | G496:C |
| O...H | 2.276 | 0.0082 | Q498:HE22 | Q42:OE1 |
| O...H | 2.429 | 0.0044 | Q498:HB2 | N501:OD1 |
| O...H | 2.857 | 0.0005 | Q498:HA | K444:O |
| O...H | 3.027 | 0.0001 | Q498:HG3 | K444:O |
| O...H | 3.828 | 0.0001 | Q498:HB3 | N501:OD1 |
| C-O | 3.938 | 0.0007 | N501:CA | Q498:O |
| C-O | 4.160 | 0.0007 | T500:C | Q498:O |
| C-O | 4.195 | 0.0005 | T500:CA | Q498:O |
| C-O | 4.478 | 0.0003 | G447:C | Q498:OE1 |
| H-C | 3.969 | 0.0002 | G447:H | Q498:CD |
| H-C | 4.075 | 0.0006 | N501:H | Q498:CA |
| H-C | 4.159 | 0.0001 | Y41:HD2 | Q498:CG |
| H-C | 4.215 | 0.0001 | N501:HB2 | Q498:C |
| H-C | 4.243 | 0.0001 | G447:HA2 | Q498:CB |
| H-C | 4.361 | 0.0002 | N501:HD21 | Q498:CA |
| H-C | 4.444 | 0.0001 | K444:H | Q498:C |
| H-C | 4.450 | 0.0001 | G446:H | Q498:CD |
| H-C | 4.476 | 0.0001 | G447:HA3 | Q498:CG |
| H-H | 3.417 | 0.0001 | Y41:HE2 | Q498:HE22 |
| H-H | 3.654 | 0.0002 | Y41:HD2 | Q498:HG2 |
| H-H | 4.140 | 0.0002 | Y41:HE2 | Q498:HG3 |
| H-H | 4.259 | 0.0001 | G447:H | Q498:HG2 |
| N...H | 4.405 | 0.0001 | G447:N | Q498:HG2 |
| N...H | 4.444 | 0.0001 | N501:N | Q498:HB3 |
| N-C | 3.765 | 0.0007 | G447:N | Q498:CD |
| N-C | 4.397 | 0.0002 | T500:N | Q498:CB |
| N-C | 4.436 | 0.0005 | N501:ND2 | Q498:CA |
| N-C | 4.479 | 0.0001 | N501:ND2 | Q498:C |
| N-O | 3.996 | 0.0003 | G447:N | Q498:OE1 |
| O...H | 2.179 | 0.0169 | N501:H | Q498:O |
| O...H | 2.381 | 0.0023 | G447:HA2 | Q498:OE1 |
| O...H | 4.136 | 0.0001 | N501:HD22 | Q498:O |
| O...H | 4.241 | 0.0001 | G447:H | Q498:OE1 |

**Table S16:** R498 with their bonding for OV interface model.

| R498 | | | | |
|---|---|---|---|---|
| H-C | 4.309 | 0.0001 | R498:HG3 | S446:CA |
| H-C | 4.312 | 0.0001 | R498:HA | S443:CA |
| H-C | 4.359 | 0.0001 | R498:H | S496:C |
| H-C | 4.371 | 0.0001 | R498:HH22 | G447:CA |
| H-C | 4.406 | 0.0002 | R498:HB3 | Y501:CE1 |
| H-C | 4.424 | 0.0002 | R498:HG3 | G447:C |
| H-C | 4.428 | 0.0002 | R498:HA | S443:C |
| H-C | 4.430 | 0.0001 | R498:HH21 | Y449:CD2 |
| H-C | 4.431 | 0.0003 | R498:HE | G447:C |
| H-C | 4.432 | 0.0003 | R498:HH11 | K353:CE |
| H-C | 4.436 | 0.0002 | R498:HH11 | D38:CG |
| H-C | 4.467 | 0.0001 | R498:HH11 | Y41:CE2 |
| H-C | 4.497 | 0.0002 | R498:HH22 | S446:C |
| H-H | 3.297 | 0.0005 | R498:HD2 | Y501:HH |
| H-H | 3.660 | 0.0004 | R498:HH12 | Y501:HH |
| H-H | 3.833 | 0.0001 | R498:HB2 | Y501:HH |
| N...H | 4.029 | 0.0001 | R498:NH1 | K353:HZ2 |
| N...H | 4.100 | 0.0002 | R498:NE | Y501:HH |
| N...H | 4.195 | 0.0007 | R498:N | N448:H |
| N...H | 4.483 | 0.0001 | R498:NE | Y41:HE2 |
| N...H | 4.496 | 0.0001 | R498:NH1 | S496:HB3 |
| N-C | 3.695 | 0.0004 | R498:NH2 | S446:C |
| N-C | 3.904 | 0.0014 | R498:NH1 | D38:CG |
| N-C | 4.211 | 0.0014 | R498:NH1 | Y501:CZ |
| N-C | 4.306 | 0.0002 | R498:N | S496:C |
| N-C | 4.332 | 0.0002 | R498:N | Y501:CE1 |
| N-C | 4.432 | 0.0006 | R498:NH1 | S496:CB |
| N-C | 4.440 | 0.0001 | R498:N | S443:CB |
| N-O | 3.957 | 0.0008 | R498:NE | Y501:OH |
| N-O | 4.056 | 0.0003 | R498:NH1 | Y449:OH |
| N-O | 4.143 | 0.0002 | R498:NH2 | D38:OD1 |
| N-O | 4.337 | 0.0001 | R498:NH1 | D38:OD2 |
| N-O | 4.410 | 0.0001 | R498:N | Y501:OH |
| O...H | 1.863 | 0.0263 | R498:HH22 | Y449:OH |
| O...H | 1.867 | 0.0525 | R498:HH21 | S446:O |
| O...H | 1.886 | 0.0345 | R498:HH12 | D38:OD1 |
| O...H | 1.948 | 0.0086 | R498:HH11 | Y501:OH |
| O...H | 2.799 | 0.0004 | R498:HG3 | K444:O |
| O...H | 2.988 | 0.0001 | R498:HA | K444:O |
| O...H | 3.254 | 0.0003 | R498:HH11 | D38:OD1 |
| O...H | 3.316 | 0.0005 | R498:HH21 | Y449:OH |
| O...H | 3.434 | 0.0004 | R498:HH12 | Y501:OH |
| O...H | 3.537 | 0.0008 | R498:HH22 | S446:O |
| O...H | 4.303 | 0.0001 | R498:HA | S443:OG |
| C-C | 4.080 | 0.0010 | S446:C | R498:CZ |
| C-C | 4.449 | 0.0002 | Y449:CZ | R498:CZ |
| C-O | 4.442 | 0.0003 | T500:CA | R498:O |
| C-O | 4.487 | 0.0001 | Y501:CA | R498:O |
| H-C | 4.066 | 0.0005 | Y501:HH | R498:CZ |
| H-C | 4.071 | 0.0002 | G447:HA3 | R498:CD |
| H-C | 4.140 | 0.0003 | Y449:HH | R498:CZ |
| H-C | 4.185 | 0.0006 | Y449:HE2 | R498:CZ |
| H-C | 4.304 | 0.0002 | G447:HA2 | R498:CB |
| H-C | 4.348 | 0.0002 | G447:H | R498:CD |
| H-C | 4.356 | 0.0001 | Q42:HG3 | R498:CZ |
| H-C | 4.357 | 0.0004 | K353:HZ1 | R498:CZ |
| H-C | 4.429 | 0.0001 | Y501:HE1 | R498:CG |
| H-C | 4.448 | 0.0001 | K353:HZ1 | R498:CD |
| H-C | 4.461 | 0.0001 | K444:H | R498:C |
| H-C | 4.482 | 0.0001 | Q42:HE21 | R498:CZ |
| H-C | 4.483 | 0.0001 | G447:HA3 | R498:CG |
| H-C | 4.493 | 0.0001 | S443:HB3 | R498:CA |
| H-H | 3.218 | 0.0002 | Y449:HH | R498:HH12 |
| H-H | 3.247 | 0.0005 | S496:HG | R498:HH11 |
| H-H | 3.359 | 0.0003 | G447:HA3 | R498:HH22 |
| H-H | 3.531 | 0.0002 | K353:HZ2 | R498:HH11 |
| H-H | 3.645 | 0.0005 | G447:HA2 | R498:HH21 |
| H-H | 3.750 | 0.0002 | K353:HZ2 | R498:HH12 |
| H-H | 3.871 | 0.0001 | K353:HZ1 | R498:HD2 |
| H-H | 3.872 | 0.0001 | G447:H | R498:HG2 |
| H-H | 3.881 | 0.0002 | S496:HB3 | R498:HH12 |
| H-H | 3.888 | 0.0001 | Q42:HG2 | R498:HH21 |
| H-H | 3.900 | 0.0001 | Q42:HE21 | R498:HH22 |
| H-H | 3.993 | 0.0001 | Q42:HG2 | R498:HH11 |
| H-H | 4.050 | 0.0002 | Y449:HH | R498:HH21 |
| H-H | 4.081 | 0.0002 | K353:HZ1 | R498:HD3 |

**Table S16:** R498 with their bonding for OV interface model.

| R498 | | | | |
|---|---|---|---|---|
| H-H | 4.087 | 0.0001 | G447:HA2 | R498:HG2 |
| H-H | 4.112 | 0.0001 | K353:HD2 | R498:HH11 |
| H-H | 4.119 | 0.0001 | S443:HA | R498:HA |
| H-H | 4.134 | 0.0001 | L45:HD23 | R498:HD3 |
| H-H | 4.229 | 0.0001 | S443:HB3 | R498:HA |
| N-C | 4.157 | 0.0006 | G447:N | R498:CZ |
| N-C | 4.158 | 0.0005 | G447:N | R498:CD |
| N-C | 4.299 | 0.0002 | Q42:NE2 | R498:CZ |
| N-C | 4.349 | 0.0001 | T500:N | R498:CB |
| N-N | 3.699 | 0.0001 | K353:NZ | R498:NH1 |
| N-N | 4.057 | 0.0014 | G447:N | R498:NH2 |
| O...H | 2.854 | 0.0016 | Y501:H | R498:O |

**Table S17:** N501 with their bonding for WT interface model.

| N501 | | | | |
|---|---|---|---|---|
| C-C | 3.795 | 0.0143 | N501:CA | G502:CA |
| C-C | 4.379 | 0.0003 | N501:CA | G502:C |
| C-O | 2.915 | 0.0007 | N501:C | T500:O |
| C-O | 4.280 | 0.0006 | N501:CB | T500:O |
| H-C | 3.713 | 0.0011 | N501:H | T500:CB |
| H-C | 3.962 | 0.0008 | N501:HB2 | T500:C |
| H-C | 4.004 | 0.0009 | N501:HA | G502:CA |
| H-C | 4.180 | 0.0003 | N501:HA | T500:CA |
| N...H | 3.172 | 0.0014 | N501:N | T500:HA |
| N...H | 3.734 | 0.0012 | N501:N | T500:HB |
| N...H | 3.955 | 0.0009 | N501:N | G502:H |
| N-C | 1.352 | 0.4225 | N501:N | T500:C |
| N-C | 3.525 | 0.0010 | N501:N | T500:CB |
| N-N | 3.625 | 0.0049 | N501:N | G502:N |
| O...H | 2.763 | 0.0017 | N501:HA | T500:O |
| O...H | 3.157 | 0.0002 | N501:HB3 | G502:O |
| O...H | 3.189 | 0.0088 | N501:H | T500:O |
| C-C | 3.734 | 0.0147 | T500:C | N501:CB |
| C-C | 3.831 | 0.0137 | T500:CA | N501:CA |
| C-C | 4.367 | 0.0004 | T500:C | N501:CG |
| C-C | 4.393 | 0.0005 | T500:CA | N501:C |
| H-C | 3.302 | 0.0120 | G502:HA3 | N501:C |
| H-C | 4.157 | 0.0005 | G502:HA2 | N501:CA |
| H-C | 4.479 | 0.0001 | T500:H | N501:CA |
| H-H | 3.399 | 0.0008 | T500:HA | N501:H |
| N...H | 3.746 | 0.0007 | G502:N | N501:HB2 |
| N-C | 1.352 | 0.4478 | G502:N | N501:C |
| N-C | 4.216 | 0.0010 | T500:N | N501:CA |
| O...H | 2.586 | 0.0008 | G502:HA2 | N501:O |
| O...H | 3.198 | 0.0081 | G502:H | N501:O |
| O...H | 3.773 | 0.0006 | G502:HA3 | N501:O |
| C-O | 3.842 | 0.0007 | N501:CG | Y505:O |
| C-O | 3.851 | 0.0015 | N501:C | K353:O |
| C-O | 3.938 | 0.0007 | N501:CA | Q498:O |
| C-O | 4.259 | 0.0005 | N501:CG | G496:O |
| H-C | 3.891 | 0.0004 | N501:HD22 | Y505:CA |
| H-C | 3.896 | 0.0004 | N501:HD21 | F497:C |
| H-C | 4.075 | 0.0006 | N501:H | Q498:CA |
| H-C | 4.100 | 0.0003 | N501:HB2 | Q506:CB |
| H-C | 4.215 | 0.0001 | N501:HB2 | Q498:C |
| H-C | 4.294 | 0.0002 | N501:HD21 | K353:CE |
| H-C | 4.361 | 0.0002 | N501:HD21 | Q498:CA |
| H-C | 4.420 | 0.0002 | N501:HB2 | Q506:CD |
| H-H | 3.725 | 0.0002 | N501:HD22 | Y505:HB2 |
| H-H | 3.739 | 0.0001 | N501:HD21 | Y505:HB3 |
| H-H | 3.824 | 0.0001 | N501:HB3 | Q506:HA |
| H-H | 4.060 | 0.0001 | N501:HB3 | Q506:HG2 |
| H-H | 4.200 | 0.0001 | N501:HB2 | Y505:HB2 |
| N...H | 4.265 | 0.0001 | N501:ND2 | Y505:HB2 |
| N...H | 4.444 | 0.0001 | N501:N | Q498:HB3 |
| N-C | 3.951 | 0.0008 | N501:ND2 | Y505:C |
| N-C | 4.264 | 0.0006 | N501:ND2 | G496:C |
| N-C | 4.264 | 0.0006 | N501:ND2 | F497:C |
| N-C | 4.436 | 0.0005 | N501:ND2 | Q498:CA |
| N-C | 4.479 | 0.0001 | N501:ND2 | Q498:C |
| O...H | 1.942 | 0.0359 | N501:HD22 | Y505:O |
| O...H | 2.179 | 0.0169 | N501:H | Q498:O |
| O...H | 2.280 | 0.0126 | N501:HD21 | G496:O |
| O...H | 3.562 | 0.0005 | N501:HD21 | Y505:O |
| O...H | 3.725 | 0.0003 | N501:HD22 | G496:O |
| O...H | 4.136 | 0.0001 | N501:HD22 | Q498:O |

**Table S18:** Y501 with their bonding for OV interface model.

| Y501 | | | | |
|---|---|---|---|---|
| C-C | 3.760 | 0.0149 | Y501:CA | G502:CA |
| C-C | 4.257 | 0.0007 | Y501:CA | G502:C |
| C-C | 4.422 | 0.0001 | Y501:CB | G502:C |
| C-O | 3.339 | 0.0003 | Y501:C | T500:O |
| C-O | 4.232 | 0.0007 | Y501:CB | T500:O |
| H-C | 3.837 | 0.0010 | Y501:H | T500:CB |
| H-C | 3.941 | 0.0010 | Y501:HA | G502:CA |
| H-C | 3.973 | 0.0007 | Y501:HB2 | T500:C |
| H-C | 4.051 | 0.0011 | Y501:HA | T500:CA |
| H-H | 3.898 | 0.0001 | Y501:HB2 | G502:H |
| N...H | 3.799 | 0.0014 | Y501:N | T500:HB |
| N...H | 3.972 | 0.0009 | Y501:N | G502:H |
| N-C | 1.357 | 0.4130 | Y501:N | T500:C |
| N-C | 3.621 | 0.0023 | Y501:N | T500:CB |
| N-N | 3.666 | 0.0050 | Y501:N | G502:N |
| N-O | 3.806 | 0.0001 | Y501:N | T500:OG1 |
| O...H | 2.525 | 0.0055 | Y501:HA | T500:O |
| O...H | 3.039 | 0.0004 | Y501:HB3 | G502:O |
| O...H | 3.172 | 0.0085 | Y501:H | T500:O |
| C-C | 3.682 | 0.0144 | T500:C | Y501:CB |
| C-C | 3.841 | 0.0132 | T500:CA | Y501:CA |
| C-C | 4.106 | 0.0004 | T500:C | Y501:CG |
| H-C | 3.297 | 0.0124 | G502:HA3 | Y501:C |
| H-C | 4.194 | 0.0003 | G502:HA2 | Y501:CA |
| H-H | 3.247 | 0.0008 | T500:HA | Y501:H |
| H-H | 3.922 | 0.0001 | T500:HB | Y501:HD1 |
| N...H | 3.711 | 0.0009 | G502:N | Y501:HB2 |
| N...H | 4.415 | 0.0001 | G502:N | Y501:HD1 |
| N-C | 1.352 | 0.4468 | G502:N | Y501:C |
| N-C | 4.212 | 0.0008 | T500:N | Y501:CA |
| N-C | 4.346 | 0.0001 | G502:N | Y501:CG |
| O...H | 3.203 | 0.0079 | G502:H | Y501:O |
| O...H | 3.779 | 0.0007 | G502:HA3 | Y501:O |
| C-O | 3.697 | 0.0008 | Y501:CE1 | Y41:OH |
| C-O | 3.935 | 0.0003 | Y501:CZ | S496:O |
| C-O | 3.937 | 0.0012 | Y501:C | K353:O |
| C-O | 4.305 | 0.0006 | Y501:CD2 | S496:O |
| C-O | 4.396 | 0.0002 | Y501:CG | H505:O |
| C-O | 4.478 | 0.0002 | Y501:CE2 | H505:O |
| C-O | 4.487 | 0.0001 | Y501:CA | R498:O |
| H-C | 2.359 | 0.0208 | Y501:HH | Y41:CD2 |
| H-C | 2.494 | 0.0140 | Y501:HH | Y41:CG |
| H-C | 3.560 | 0.0007 | Y501:HD1 | Y41:CZ |
| H-C | 3.657 | 0.0002 | Y501:HD1 | Y41:CE1 |
| H-C | 3.783 | 0.0006 | Y501:HE2 | K353:CE |
| H-C | 3.887 | 0.0002 | Y501:HH | K353:CE |
| H-C | 3.901 | 0.0001 | Y501:HE2 | F497:C |
| H-C | 4.066 | 0.0005 | Y501:HH | R498:CZ |
| H-C | 4.086 | 0.0004 | Y501:HB2 | Q506:CB |
| H-C | 4.191 | 0.0002 | Y501:HD2 | H505:CA |
| H-C | 4.256 | 0.0003 | Y501:HB2 | Q506:CD |
| H-C | 4.386 | 0.0004 | Y501:HD1 | Y41:CE2 |
| H-C | 4.400 | 0.0001 | Y501:HB3 | K353:C |
| H-C | 4.412 | 0.0002 | Y501:HB3 | H505:CG |
| H-C | 4.429 | 0.0001 | Y501:HE1 | R498:CG |
| H-C | 4.434 | 0.0001 | Y501:HA | D355:CA |
| H-C | 4.437 | 0.0001 | Y501:HA | D355:CG |
| H-C | 4.482 | 0.0001 | Y501:HE1 | Y41:CB |
| H-H | 3.644 | 0.0001 | Y501:HD2 | H505:HB2 |
| H-H | 3.737 | 0.0001 | Y501:HB2 | Q506:HE22 |
| H-H | 3.880 | 0.0002 | Y501:HB2 | H505:HB2 |
| H-H | 4.038 | 0.0001 | Y501:HB3 | Q506:HG2 |
| H-H | 4.054 | 0.0001 | Y501:H | Q506:HG3 |
| N...H | 4.285 | 0.0001 | Y501:N | D355:HB2 |
| O...H | 2.213 | 0.0016 | Y501:HE2 | S496:O |
| O...H | 2.854 | 0.0016 | Y501:H | R498:O |
| O...H | 4.358 | 0.0002 | Y501:HD2 | S496:O |
| O...H | 4.384 | 0.0001 | Y501:HE2 | H505:O |
| C-C | 3.894 | 0.0003 | Y41:CG | Y501:CE1 |
| C-C | 4.033 | 0.0013 | Y41:CD2 | Y501:CZ |
| C-C | 4.082 | 0.0009 | K353:CE | Y501:CZ |
| C-C | 4.114 | 0.0007 | K353:CE | Y501:CE2 |
| C-C | 4.123 | 0.0004 | Y41:CG | Y501:CZ |
| C-C | 4.133 | 0.0010 | Y41:CE2 | Y501:CZ |
| C-C | 4.151 | 0.0002 | F497:C | Y501:CE2 |
| C-C | 4.255 | 0.0014 | S496:C | Y501:CE2 |

**Table S17:** N501 with their bonding for WT interface model.

| N501 | | | | |
|---|---|---|---|---|
| C-C | 4.318 | 0.0004 | Q498:CA | N501:CG |
| C-O | 3.697 | 0.0008 | Q506:CD | N501:O |
| C-O | 3.989 | 0.0002 | Q498:CA | N501:OD1 |
| H-C | 3.914 | 0.0007 | F497:HA | N501:CG |
| H-C | 4.074 | 0.0004 | Q506:HE22 | N501:CA |
| H-C | 4.272 | 0.0001 | Y505:HB2 | N501:CG |
| H-C | 4.281 | 0.0001 | Q506:HG2 | N501:CA |
| H-C | 4.402 | 0.0001 | Y505:HB2 | N501:C |
| H-C | 4.429 | 0.0001 | Q506:HA | N501:CG |
| H-C | 4.458 | 0.0002 | Q498:HB2 | N501:CB |
| H-C | 4.475 | 0.0001 | Y505:HB2 | N501:CA |
| H-H | 3.210 | 0.0005 | Q498:H | N501:HD22 |
| H-H | 3.359 | 0.0001 | F497:HD1 | N501:HD21 |
| H-H | 3.410 | 0.0001 | K353:HD3 | N501:HD22 |
| N-C | 4.064 | 0.0026 | Q506:NE2 | N501:C |
| N-C | 4.220 | 0.0007 | V503:N | N501:C |
| N-N | 4.488 | 0.0005 | P499:N | N501:N |
| O...H | 1.869 | 0.0394 | Q506:HE22 | N501:O |
| O...H | 2.429 | 0.0044 | Q498:HB2 | N501:OD1 |
| O...H | 3.553 | 0.0005 | Q506:HE21 | N501:O |
| O...H | 3.828 | 0.0001 | Q498:HB3 | N501:OD1 |
| O...H | 4.252 | 0.0001 | V503:HA | N501:O |

**Table S18:** Y501 with their bonding for OV interface model.

| Y501 | | | | |
|---|---|---|---|---|
| C-C | 4.300 | 0.0013 | Y41:CZ | Y501:CZ |
| C-C | 4.363 | 0.0006 | Y41:CE1 | Y501:CZ |
| C-C | 4.415 | 0.0001 | K353:CB | Y501:CZ |
| C-C | 4.435 | 0.0008 | Y41:CE2 | Y501:CD1 |
| C-C | 4.455 | 0.0001 | R498:CG | Y501:CZ |
| C-C | 4.464 | 0.0002 | R498:CA | Y501:CZ |
| C-C | 4.478 | 0.0009 | K353:CG | Y501:CZ |
| C-C | 4.481 | 0.0005 | R498:CA | Y501:CE2 |
| C-O | 3.573 | 0.0019 | K353:CE | Y501:OH |
| C-O | 3.786 | 0.0009 | Q506:CD | Y501:O |
| C-O | 3.905 | 0.0014 | R498:CZ | Y501:OH |
| C-O | 4.359 | 0.0007 | R498:CG | Y501:OH |
| C-O | 4.389 | 0.0008 | Y41:CZ | Y501:OH |
| C-O | 4.426 | 0.0002 | R498:CB | Y501:OH |
| C-O | 4.471 | 0.0007 | Y41:CE1 | Y501:OH |
| H-C | 3.688 | 0.0003 | K353:HZ2 | Y501:CZ |
| H-C | 3.883 | 0.0021 | K353:HZ1 | Y501:CE1 |
| H-C | 3.994 | 0.0004 | F497:HA | Y501:CZ |
| H-C | 4.045 | 0.0001 | H505:HB2 | Y501:CG |
| H-C | 4.066 | 0.0001 | H505:HB2 | Y501:CD2 |
| H-C | 4.116 | 0.0005 | R498:HH11 | Y501:CE1 |
| H-C | 4.189 | 0.0001 | D355:HB2 | Y501:C |
| H-C | 4.205 | 0.0001 | Q506:HG2 | Y501:CA |
| H-C | 4.259 | 0.0001 | Q506:HG2 | Y501:C |
| H-C | 4.323 | 0.0003 | K353:HZ1 | Y501:CD2 |
| H-C | 4.365 | 0.0003 | K353:HZ3 | Y501:CZ |
| H-C | 4.367 | 0.0001 | H505:HB2 | Y501:C |
| H-C | 4.406 | 0.0002 | R498:HB3 | Y501:CE1 |
| H-C | 4.436 | 0.0002 | H505:HB2 | Y501:CA |
| H-C | 4.461 | 0.0002 | Y41:HD2 | Y501:CZ |
| H-H | 3.297 | 0.0005 | R498:HD2 | Y501:HH |
| H-H | 3.660 | 0.0004 | R498:HH12 | Y501:HH |
| H-H | 3.717 | 0.0001 | F497:HD1 | Y501:HE2 |
| H-H | 3.792 | 0.0001 | K353:HZ3 | Y501:HH |
| H-H | 3.833 | 0.0001 | R498:HB2 | Y501:HH |
| H-H | 3.932 | 0.0002 | K353:HZ2 | Y501:HH |
| H-H | 3.934 | 0.0004 | K353:HZ3 | Y501:HE2 |
| H-H | 3.952 | 0.0001 | K353:HE3 | Y501:HH |
| H-H | 4.061 | 0.0001 | K353:HD3 | Y501:HH |
| H-H | 4.309 | 0.0001 | D355:HB3 | Y501:HA |
| N...H | 4.100 | 0.0002 | R498:NE | Y501:HH |
| N...H | 4.355 | 0.0001 | H505:N | Y501:HB3 |
| N-C | 3.589 | 0.0011 | K353:NZ | Y501:CZ |
| N-C | 3.996 | 0.0021 | Q506:NE2 | Y501:C |
| N-C | 4.060 | 0.0010 | V503:N | Y501:C |
| N-C | 4.211 | 0.0014 | R498:NH1 | Y501:CZ |
| N-C | 4.332 | 0.0002 | R498:N | Y501:CE1 |
| N-O | 3.957 | 0.0008 | R498:NE | Y501:OH |
| N-O | 4.410 | 0.0001 | R498:N | Y501:OH |
| O...H | 1.782 | 0.0518 | K353:HZ1 | Y501:OH |
| O...H | 1.864 | 0.0425 | Q506:HE22 | Y501:O |
| O...H | 1.948 | 0.0086 | R498:HH11 | Y501:OH |
| O...H | 3.434 | 0.0004 | R498:HH12 | Y501:OH |
| O...H | 3.495 | 0.0003 | Q506:HE21 | Y501:O |
| O...H | 3.575 | 0.0004 | K353:HD3 | Y501:OH |
| O...H | 3.882 | 0.0001 | V503:H | Y501:O |
| O...H | 3.890 | 0.0002 | K353:HE3 | Y501:OH |
| O...H | 3.985 | 0.0001 | V503:HA | Y501:O |
| O...H | 4.244 | 0.0001 | Y41:HB2 | Y501:OH |

**Table S19:** Y505 with their bonding for WT interface model.

| Y505 | | | | |
|---|---|---|---|---|
| C-C | 3.707 | 0.0128 | Y505:C | Q506:CB |
| C-C | 3.841 | 0.0130 | Y505:CA | Q506:CA |
| C-O | 3.915 | 0.0008 | Y505:CB | G504:O |
| C-O | 4.019 | 0.0004 | Y505:C | G504:O |
| H-C | 3.787 | 0.0001 | Y505:HB2 | G504:C |
| H-C | 4.047 | 0.0017 | Y505:HA | G504:CA |
| H-C | 4.140 | 0.0002 | Y505:H | Q506:CA |
| H-H | 3.087 | 0.0002 | Y505:HA | Q506:H |
| N...H | 3.123 | 0.0001 | Y505:N | G504:HA3 |
| N...H | 3.208 | 0.0021 | Y505:N | G504:HA2 |
| N-C | 1.360 | 0.4017 | Y505:N | G504:C |
| N-C | 4.176 | 0.0014 | Y505:N | Q506:CA |
| O...H | 2.398 | 0.0075 | Y505:HA | G504:O |
| O...H | 3.202 | 0.0083 | Y505:H | G504:O |
| C-C | 3.520 | 0.0046 | G504:C | Y505:CB |
| C-C | 3.587 | 0.0077 | G504:C | Y505:C |
| C-C | 3.853 | 0.0130 | G504:CA | Y505:CA |
| C-O | 3.459 | 0.0011 | Q506:C | Y505:O |
| C-O | 4.214 | 0.0006 | Q506:CB | Y505:O |
| H-C | 3.927 | 0.0016 | Q506:H | Y505:CB |
| H-C | 3.946 | 0.0008 | Q506:HB2 | Y505:C |
| H-C | 4.084 | 0.0011 | Q506:HA | Y505:CA |
| H-C | 4.397 | 0.0001 | Q506:HG2 | Y505:C |
| H-H | 3.286 | 0.0004 | G504:HA3 | Y505:H |
| H-H | 3.442 | 0.0007 | G504:HA2 | Y505:H |
| N...H | 3.800 | 0.0015 | Q506:N | Y505:HB3 |
| N...H | 3.828 | 0.0004 | Q506:N | Y505:HB2 |
| N-C | 1.346 | 0.4603 | Q506:N | Y505:C |
| N-C | 3.661 | 0.0034 | Q506:N | Y505:CB |
| N-C | 4.269 | 0.0007 | G504:N | Y505:CA |
| O...H | 2.444 | 0.0034 | Q506:HA | Y505:O |
| O...H | 3.171 | 0.0085 | Q506:H | Y505:O |
| C-O | 3.552 | 0.0014 | Y505:CA | G502:O |
| C-O | 4.187 | 0.0001 | Y505:CG | K353:O |
| H-C | 4.016 | 0.0010 | Y505:H | G502:CA |
| H-C | 4.169 | 0.0001 | Y505:HE1 | K353:C |
| H-C | 4.261 | 0.0001 | Y505:HD1 | K353:CB |
| H-C | 4.272 | 0.0001 | Y505:HB2 | N501:CG |
| H-C | 4.402 | 0.0001 | Y505:HB2 | N501:C |
| H-C | 4.475 | 0.0001 | Y505:HB2 | N501:CA |
| O...H | 1.901 | 0.0335 | Y505:H | G502:O |
| N-C | 3.861 | 0.0006 | Y505:N | G502:C |
| C-C | 4.303 | 0.0002 | K353:CB | Y505:CD1 |
| C-C | 4.348 | 0.0002 | K353:C | Y505:CG |
| C-C | 4.488 | 0.0001 | K353:CA | Y505:CB |
| C-O | 3.842 | 0.0007 | N501:CG | Y505:O |
| H-C | 3.891 | 0.0004 | N501:HD22 | Y505:CA |
| H-C | 3.965 | 0.0002 | K353:HB3 | Y505:CD1 |
| H-C | 4.300 | 0.0002 | K353:H | Y505:CE1 |
| H-C | 4.480 | 0.0001 | P507:HD3 | Y505:C |
| H-H | 3.725 | 0.0002 | N501:HD22 | Y505:HB2 |
| H-H | 3.739 | 0.0001 | N501:HD21 | Y505:HB3 |
| H-H | 3.871 | 0.0001 | K353:HB3 | Y505:HD1 |
| H-H | 4.200 | 0.0001 | N501:HB2 | Y505:HB2 |
| N...H | 4.265 | 0.0001 | N501:ND2 | Y505:HB2 |
| N-C | 3.951 | 0.0008 | N501:ND2 | Y505:C |
| N-C | 4.281 | 0.0004 | P507:N | Y505:C |
| N-N | 4.396 | 0.0005 | V503:N | Y505:N |
| O...H | 1.942 | 0.0359 | N501:HD22 | Y505:O |
| O...H | 3.562 | 0.0005 | N501:HD21 | Y505:O |
| O...H | 3.842 | 0.0001 | P507:HD3 | Y505:O |

**Table S20:** H505 with their bonding for OV interface

| H505 | | | | |
|---|---|---|---|---|
| C-C | 3.719 | 0.0133 | H505:C | Q506:CB |
| C-C | 3.840 | 0.0133 | H505:CA | Q506:CA |
| C-O | 3.933 | 0.0004 | H505:C | G504:O |
| C-O | 4.020 | 0.0008 | H505:CB | G504:O |
| H-C | 3.861 | 0.0001 | H505:HB2 | G504:C |
| H-C | 4.042 | 0.0016 | H505:HA | G504:CA |
| H-C | 4.217 | 0.0002 | H505:H | Q506:CA |
| H-H | 3.181 | 0.0005 | H505:HA | Q506:H |
| H-H | 3.833 | 0.0001 | H505:HB2 | Q506:H |
| N...H | 3.233 | 0.0025 | H505:N | G504:HA2 |
| N...H | 4.185 | 0.0001 | H505:ND1 | G504:HA3 |
| N-C | 1.359 | 0.4009 | H505:N | G504:C |
| N-C | 4.162 | 0.0014 | H505:N | Q506:CA |
| O...H | 2.390 | 0.0070 | H505:HA | G504:O |
| O...H | 3.198 | 0.0083 | H505:H | G504:O |
| C-C | 3.556 | 0.0063 | G504:C | H505:C |
| C-C | 3.568 | 0.0066 | G504:C | H505:CB |
| C-C | 3.853 | 0.0130 | G504:CA | H505:CA |
| C-O | 3.390 | 0.0009 | Q506:C | H505:O |
| C-O | 4.251 | 0.0006 | Q506:CB | H505:O |
| H-C | 3.833 | 0.0017 | Q506:H | H505:CB |
| H-C | 3.946 | 0.0008 | Q506:HB2 | H505:C |
| H-C | 4.084 | 0.0010 | Q506:HA | H505:CA |
| H-C | 4.400 | 0.0001 | Q506:HG2 | H505:C |
| H-H | 3.222 | 0.0004 | G504:HA3 | H505:H |
| H-H | 3.484 | 0.0007 | G504:HA2 | H505:H |
| N...H | 3.655 | 0.0005 | Q506:N | H505:HB2 |
| N...H | 3.750 | 0.0016 | Q506:N | H505:HB3 |
| N-C | 1.351 | 0.4335 | Q506:N | H505:C |
| N-C | 3.587 | 0.0027 | Q506:N | H505:CB |
| N-C | 4.288 | 0.0006 | G504:N | H505:CA |
| O...H | 2.496 | 0.0041 | Q506:HA | H505:O |
| O...H | 3.174 | 0.0086 | Q506:H | H505:O |
| C-O | 3.553 | 0.0005 | H505:CD2 | E37:OE1 |
| C-O | 3.653 | 0.0013 | H505:CA | G502:O |
| C-O | 3.661 | 0.0014 | H505:CE1 | E37:OE1 |
| H-C | 3.984 | 0.0007 | H505:H | G502:CA |
| H-C | 4.045 | 0.0001 | H505:HB2 | Y501:CG |
| H-C | 4.066 | 0.0001 | H505:HB2 | Y501:CD2 |
| H-C | 4.100 | 0.0003 | H505:HE2 | K353:CB |
| H-C | 4.367 | 0.0001 | H505:HB2 | Y501:C |
| H-C | 4.404 | 0.0001 | H505:HB3 | K353:CG |
| H-C | 4.436 | 0.0002 | H505:HB2 | Y501:CA |
| N...H | 3.965 | 0.0001 | H505:NE2 | K353:HB3 |
| N...H | 4.355 | 0.0001 | H505:N | Y501:HB3 |
| N-C | 3.918 | 0.0036 | H505:NE2 | E37:CD |
| N-C | 4.071 | 0.0005 | H505:NE2 | K353:CB |
| N-C | 4.417 | 0.0002 | H505:NE2 | K353:C |
| O...H | 1.767 | 0.0460 | H505:HE2 | E37:OE1 |
| O...H | 1.981 | 0.0258 | H505:H | G502:O |
| O...H | 3.626 | 0.0004 | H505:HD2 | E37:OE1 |
| O...H | 3.842 | 0.0002 | H505:HE1 | E37:OE1 |
| O...H | 4.020 | 0.0010 | H505:HE2 | E37:OE2 |
| C-C | 4.420 | 0.0002 | K353:C | H505:CD2 |
| C-O | 4.396 | 0.0002 | Y501:CG | H505:O |
| C-O | 4.478 | 0.0002 | Y501:CE2 | H505:O |
| H-C | 3.845 | 0.0002 | K353:HG2 | H505:CG |
| H-C | 4.072 | 0.0001 | K353:HA | H505:CB |
| H-C | 4.191 | 0.0002 | Y501:HD2 | H505:CA |
| H-C | 4.202 | 0.0001 | K353:HD3 | H505:CG |
| H-C | 4.412 | 0.0002 | Y501:HB3 | H505:CG |
| H-C | 4.482 | 0.0001 | K353:H | H505:CE1 |
| H-H | 3.644 | 0.0001 | Y501:HD2 | H505:HB2 |
| H-H | 3.792 | 0.0001 | E37:HG3 | H505:HE2 |
| H-H | 3.880 | 0.0002 | Y501:HB2 | H505:HB2 |
| H-H | 3.942 | 0.0001 | K353:HA | H505:HB2 |
| H-H | 4.108 | 0.0001 | K353:HG3 | H505:HD2 |
| H-H | 4.230 | 0.0001 | K353:HG2 | H505:HB3 |
| N-C | 4.286 | 0.0003 | P507:N | H505:C |
| N-N | 4.348 | 0.0002 | K353:N | H505:NE2 |
| N-N | 4.432 | 0.0004 | V503:N | H505:N |
| O...H | 3.933 | 0.0001 | P507:HD3 | H505:O |
| O...H | 4.384 | 0.0001 | Y501:HE2 | H505:O |

**References:**


1. Package, V.-V.A.i.S.; Available from: https://www.vasp.at/.
2. Perdew, J.P., K. Burke, and M. Ernzerhof, Generalized gradient approximation made simple. *Phys. Rev. Lett.*, **1996**. 77(18): p. 3865.
3. Ching, W.-Y. and P. Rulis, Electronic Structure Methods for Complex Materials: The orthogonalized linear combination of atomic orbitals. **2012**: Oxford University Press.
4. Mulliken, R.S., Electronic population analysis on LCAO–MO molecular wave functions. I. *J. Chem. Phys.*, **1955**. 23(10): p. 1833-1840.
5. Mulliken, R., Electronic population analysis on LCAO–MO molecular wave functions. II. Overlap populations, bond orders, and covalent bond energies. *J. Chem. Phys.*, **1955**. 23(10): p. 1841-1846.
